# A Quality Measure for Repeating Multiple-Unit Spike Patterns

**DOI:** 10.64898/2026.01.31.702754

**Authors:** Günther Palm, Monica Paoletti, Junji Ito, Alessandra Stella, Sonja Grün

## Abstract

We propose a quality measure for spatio-temporal spike patterns (STPs) in multiple-neuron recordings. In such recordings, repeating STPs or pattern repetitions (PRs) are often found, with many of these generated by chance. To rule those out, statistical tests have been developed to discriminate the unlikely from the more likely PRs. This statistical problem is complicated by the fact that there are several obvious quality criteria for a PR, such as the size (the number of spikes) of the pattern and the number of its occurrences. Here, we propose a canonical way of combining several criteria (which we collect in the so-called signature of the pattern) into a single quality measure, based on the ‘unlikeliness’ of the pattern. This measure is defined mathematically, and a formula for its computation is derived for stationary spike trains. It can be used to compare PRs. Since spike trains are not stationary in practice, we discuss, for two experimental data sets, how well the stationary formula correlates with the defined quality measure as determined from simulations. The results encourage the use of the stationary formula or also some simpler, related formulas as ‘proxies’ for the quality, for the comparison of PRs and also for statistical tests that avoid the multiple testing problem incurred by using several quality criteria. Based on our results, we propose a few test statistics, i.e., random variables on the space of multi-unit spike trains with an appropriate null-hypothesis distribution, to evaluate STPs with less computational and sampling efforts.

## 1 Introduction

A basic goal of neurophysiology is to understand the internal representation of the world outside in the neuronal activation in the animal’s brain. To study this representation, spike trains of single neurons were recorded and correlated to experimental sensory stimuli, primarily in sensory brain areas (Barlow 1961; Perkel and Bullock 1967). To answer the computational question of how these neural responses can be generated in the network of neurons forming the brain, one has to study the interactions between these neurons. Correlated spiking activity was suggested as the physiological indication of neuronal interaction during this computational process, initially by George Gerstein (Aertsen et al. 2023). Most analyses were restricted to pairwise correlations, even when recording from multiple neurons became possible, because the sheer number of correlograms to be analyzed increases almost exponentially with increasing order, and they also become much harder to display and interpret (Palm and Pöpel (1985); Staude et al. (2010a,Staude et al. 2010b); Gerstein et al. (2012), and in Aertsen et al. (2023); Grün and Rotter (2010)); only a few went up to third order, e.g., Perkel et al. (1975); Martignon et al. (1995); Tetko and Villa (2001); Prut et al. (1998); Staude et al. (2010a). However, if one considers only positive interactions among (sparsely occurring) spikes, it may be possible to analyze the interaction of larger groups of neurons in multiple-neuron spike patterns, which are typically assumed in models of brain function (Hebb 1949; Palm 1982; Abeles 1982). These patterns could be synchronous spike patterns or spike patterns with specific time delays between the individual spikes. Clearly, such patterns have to occur repeatedly in multiple-unit recordings in order to be of interest; they are simply called (spatio-temporal) spike patterns.

Identifying such patterns requires a systematic search for all possible spike patterns in multiple-unit recordings. This will typically result in a huge number of patterns (millions), most of which are likely not interesting, because they are either too small or not repeated sufficiently often. In addition, such searches are computationally very expensive. Some early methods found repeating patterns in multiple-unit recordings of relatively small numbers of neurons (Abeles et al. 1993; Abeles and Gat 2001; Shmiel et al. 2006; Prut et al. 1998; Nádasdy et al. 1999; Shimazaki et al. 2012). Some methods have been developed to perform these computations on a larger number of neurons and more efficiently (Quaglio et al. (2018) and citations therein or Schrader et al. (2008); Torre et al. (2013, 2016b)), usually based on advanced computer science methods like frequent itemset mining (FIM) (Picado-Muiño et al. 2013). Still, one often ends up with too many patterns to analyze in detail.

This type of research has led to the finding of many repeating spike patterns in various animals and cortical areas that are related to behavior and learning (Vaadia et al. 1995; Prut et al. 1998; Riehle et al. 1997; Ikegaya et al. 2004; Kilavik et al. 2009; Fellous et al. 2004; Srivastava et al. 2017; Fiebelkorn and Kastner 2020), but also to intense discussions about the general physiological and statistical relevance of such patterns (Baker and Lemon 2000; Kass et al. 2014, Kass et al. 2005). In particular, statistical tests have been developed to show that there are more repeated spike patterns and they are repeated more often than one would expect just by chance. More concretely, the focus on the sheer number of repeating ‘spatio-temporal’ patterns has sometimes turned out to be non-conclusive, because without strong restrictions on pattern size and number of repetitions, too many patterns occur by chance (Baker and Lemon 2000), while the idea to determine the significance of individual patterns is also problematic, due to the vast number of potential patterns which incurs a vast multiple testing problem (Kirsch et al. 2012). Therefore, in particular in the SPADE project (Torre et al. 2016b; Quaglio et al. 2017; Stella et al. 2019, 2022) the focus shifted to testing the unexpectedness of classes of pattern repetition events that are highly unlikely, in particular due to a large number of repetitions (Torre et al. 2016b; Quaglio et al. 2017; Stella et al. 2019, 2022). One problem with these tests is that they still are too strict, because several criteria can make a pattern repetition unexpected and could serve as test statistics, and since it is not obvious how to combine them, one still has to resort to multiple testing procedures, which result in very small numbers of significant patterns (Stella 2022), even with multiple testing corrections that control the false discovery rate (Benjamini and Hochberg 1995).

Here, we show a way out of this dilemma: we define a new measure for the ‘quality’ of a pattern repetition that can be used to compare different pattern repetition events, and we suggest using it (or a simpler approximation to it) to define the test variable for a statistical test. To be sure, in today’s research, such a statistical test is not really used to reject the hypothesis that all spike patterns can be attributed just to chance, since there is ample evidence for the existence and behavioral relevance of such patterns (citations were given above). In practice, significance probabilities may rather be used to compare different pattern repetitions,or to set a threshold, beyond which individual pattern repetitions can be considered as sufficiently interesting for further analysis. The ultimate argument for the relevance of spike patterns will, of course, always be their relation to the recorded behavior, i.e., their repeated occurrence at particular moments in time during the experimental protocol. Here, our aim is just to simplify the comparison and statistical analysis of repeating precisely timed spike patterns.

Before we introduce our statistical model, let us briefly mention the quality criteria that can be used to compare repeating spike patterns. The first two most obvious quality criteria are the **size** of the pattern (i.e., the number of spikes), and how often it occurs, i.e., its **repetition**. In reality, a pattern is never repeated exactly. Therefore, the **precision** of pattern repetitions can also be considered as a quality criterion. But temporal imprecision may not be the only inexactness to be tolerated: one or two spikes may be missing, the order may be changed, and so on. Such considerations can result in the introduction of several more numerical quality criteria for pattern repetitions, and the relative weighting of these criteria will certainly be a matter of taste. Much of this is reflected in the discussion of various definitions of a similarity or distance measure between two spike trains (van Rossum et al. 2000; Houghton and Sen 2008; Victor and Purpura 1997; Humphries 2011), which can also be extended to multiple-unit spike trains (Sotomayor-Gómez et al. 2023) and used as a measure for the similarity of repeated STPs.

Thus, the first goal of this paper is to **propose a canonical way of combining several intuitive quality criteria into one quality measure**. It is based on the argument that each quality criterion should serve the purpose of reducing the set of potential patterns and pattern repetitions and that the agreement on several criteria will reduce it even further. Therefore, the combined quality measure can be based on **the number of potential patterns that fulfill all accepted quality criteria**. So, our mathematical goal is to define and estimate this number mathematically, which we call **ubiquity**. Such an estimate can also be very useful in the practical design of algorithms for extensive pattern search, because it provides a reasonable estimate of the number of chance patterns we have to expect that fulfill a given set of quality criteria and therefore the amount of computational resources required to handle them. Since larger ubiquity implies lower quality, we define the **quality** as the negative logarithm of ubiquity. Here we show the line of argument for such a computation for a particular collection of quality criteria, which we define in Section 3.1. Based on this quality measure, we develop statistical tests for the surprisingness of a pattern repetition event, or of the whole multiple-unit spike train in terms of pattern repetitions. In addition, we present an important new quality criterion here, namely the so-called **novelty** of a repeated pattern, which is used to represent the firing rates of the neurons that fire repeatedly in the pattern. The firing rates should be considered, because the expected number of repetitions of a pattern will be larger if the participating neurons have larger firing rates.

Since neurons usually do not respond stereotypically to repetitions of the same experimental condition, but can show a considerable variability (Softky and Koch 1993, Softky and Koch 1992; Knoblauch and Palm 2004), their response spike-trains are typically modeled as stochastic processes, in particular as point processes, where the ‘point’ events correspond to the spikes (Perkel et al. 1967; Brown et al. 2004; Grün and Rotter 2010; Cox and Isham 1980; Cox and Lewis 1966). This does not necessarily mean that the ‘reason’ behind this apparent stochasticity is some ‘intrinsic’ physical randomness occurring somewhere inside the neurons or synapses (Mainen and Sejnowski 1995). Instead, the use of a stochastic description may reflect the interaction of the observed neuron(s) with numerous unobserved ones, which follow a flow of activation through the observed cortical area or essentially the whole brain of the animal (Knoblauch and Palm 2004). In this view, the brain is described as a complex, deterministic, but high-dimensional dynamical system, and the recorded multiple-neuron activity is viewed as a low-dimensional projection from it (Palm 1982, 2022; Gerstner et al. 2014; Freeman 2000). The stochastic description of neuronal spike trains comes with the use of methods from probability, stochastic processes, and also information theory, often described as the search for the ‘neural code’, i.e., the representation of a particular aspect of the external situation in the population activity of all neurons in a particular area of the brain or the cortex. Here we will use these stochastic methods to define our statistical ‘null hypotheses’, the **multiple-unit spike train (MUST)**, which also serves as the framework to formally define the quality criteria that we use. Our null hypothesis essentially assumes that each neuron follows the experimental conditions with a variable firing rate, but independent of the other neurons, i.e., we allow for non-stationary spike trains with covariation due to the sensory input, but without interactions between different neurons. And we use the quality of pattern repetitions as evidence against this null hypothesis.

In the next section, we define the statistical framework, i.e., the multiple-unit spike train as a statistical process, the spatio-temporal pattern (STP), the pattern repetition event and its *novelty*, the pre-order given by several quality criteria, and finally the expected number of relevant patterns and the *quality* measure. In Section 3.1, we show how to calculate the values of these quantities for specific patterns under the assumption of a stationary process. Since the assumption of stationarity is typically unrealistic in neurophysiology, we then compare in Section 4 the calculated ubiquity of patterns (assuming stationarity) and the ubiquity obtained from simulations of non-stationary spike trains for different typical experimental situations. Based on these comparisons, we will argue that with a bit of caution, the calculated ‘stationary quality’ may still be useful as a reasonable way of combining various quality criteria into one. We will also show some other combinations of our quality criteria for comparison. In Section 5, we discuss in more detail how the MUST and the quality calculations can be generated for a concrete multiple-unit data set and how the statistical quality test can be performed.

## 2 Defining novelty and quality of a repeating pattern

In this section, we define the mathematical framework for our concept of spatio-temporal spike-patterns or STPs and their quality criteria, leading up to the definition of the *ubiquity* of a pattern with a certain quality, which is the expected number of patterns that fulfill particular quality criteria. Then the negative logarithm of the ubiquity is defined as the numerical *quality* of the pattern.

Usually, spike trains are modeled as point processes, but here we simplify this slightly by using the assumption of discrete time. This is reasonable since time is discretized anyway at the resolution of the experimental recording, and in many cases, time bins are used, which allow for a coarse distinction of spike times that simplifies the evaluation. The optimal choice of the time resolution and the method of binning is itself a complicated issue that we don’t consider here.

In the discrete-time framework, multiple-unit spike trains are just represented as a collection of random variables. Also, other entities will be described as random variables, and we will use the convention that they are denoted by capital letters, whereas their individual values are usually denoted by the respective lowercase letters.

### Definition 1

(multiple-neuron spike train) Let *N* be the number of neurons and *T* be the total time of observation in bins. Let 𝒩 *=* {1,…, *N* } and 𝒯 *=* {0,…, *T* − 1} be the set of neuron and time indices, respectively. Let Ξ *=* {0, 1}^𝒩 *×*𝒯^ be the space of all possible multiple-unit spike trains 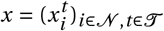, where 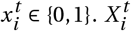 is the corresponding binary random variable that represents the neural activity of neuron *i* at time *t* and

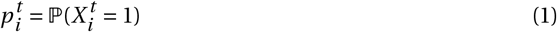

is the probability of neuron *i* to emit a spike at time *t* . The collection 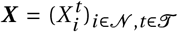 of these random variables is considered as a stochastic process that describes a **multiple-unit spike-train (MUST)**. An element *x* ∈ Ξ is also called a multiple-unit spike-train (MUST).

Technically, Ξ is a finite (though very large) probability space on which all random variables considered in this paper are defined. Usually, we will assume that all these random variables that describe the MUST are independent. This is, of course, not what neuroscientists really believe, but it can serve as a first null-hypothesis that expresses the idea of completely random generation of patterns. Considering this idea of a null-hypothesis in a little more detail, it is often argued that these spike patterns are due to specific interactions between individual neurons beyond their common reaction to the stimulus or behavioral situation. In this case, one may consider the assumption of independence of spike events on different neurons, but perhaps not of spikes of the same neuron at subsequent times, because there are several biophysical mechanisms in single neurons, like refractoriness or burstiness (Harris et al. 2001; Avila-Akerberg and Chacron 2011), which should be taken into account even in a null-hypothesis. However, these effects are typically limited to a short time scale up to perhaps 10 or 20 ms. Here, it may be assumed in a null hypothesis for simplicity that spike events at longer delays are (approximately) independent. If one is not particularly interested in Morse-code patterns generated by single neurons, one may largely avoid this discussion by simply excluding subsequent spikes on the same neuron from the definition of an STP; in addition, one could also ask for a minimal time delay between subsequent occurrences of an STP.

We do not require *stationarity* of the spike trains for our definitions; however, for the mathematical proofs in the next section, we will have to assume it. Stationarity means that the spike probabilities are independent of time *t*, i.e., 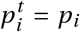 only depends on the neuron *i* . Again, this assumption is problematic from the experimental point of view, but it drastically reduces the number of model parameters and makes the calculations feasible.

In practice, experimental spike trains are usually not stationary, and it is also not obvious how to determine the ‘instantaneous’ firing probabilities 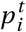 that we need for our definition of the MUST. Here we have to rely on averaging the spike counts across a suitable time window, for example, as in Baker and Lemon (2000), or estimate the probabilities based on inter-spike intervals, as described in Section 3.3.

### Definition 2

(spike pattern and STP) Let *W* be the size of the time window that is used in the definition and detection of patterns. Let 𝒲 *=* {0,…, *W* − 1} be the set of time indices within a window. A **spike pattern** is a subset *P* of 𝒩 *×* 𝒲 . The quantity *z =* |*P*| is called the **size** of the pattern. Here we require that *z* ≥ 2, and we usually consider only spike patterns with *z* ≥ 3.

We consider the sets

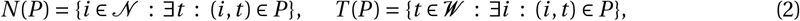

which lists the neurons and the time points of the pattern. The **duration** of the pattern *P* is

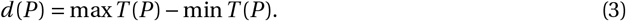

A spike pattern *P* ⊆ 𝒩 *×*𝒲 such that

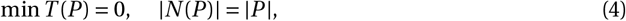

is called a **spatio-temporal pattern (STP)**. The set of all STPs is called 𝒫 .

The concept of an STP is illustrated in Fig. 1.

**Fig. 1:**
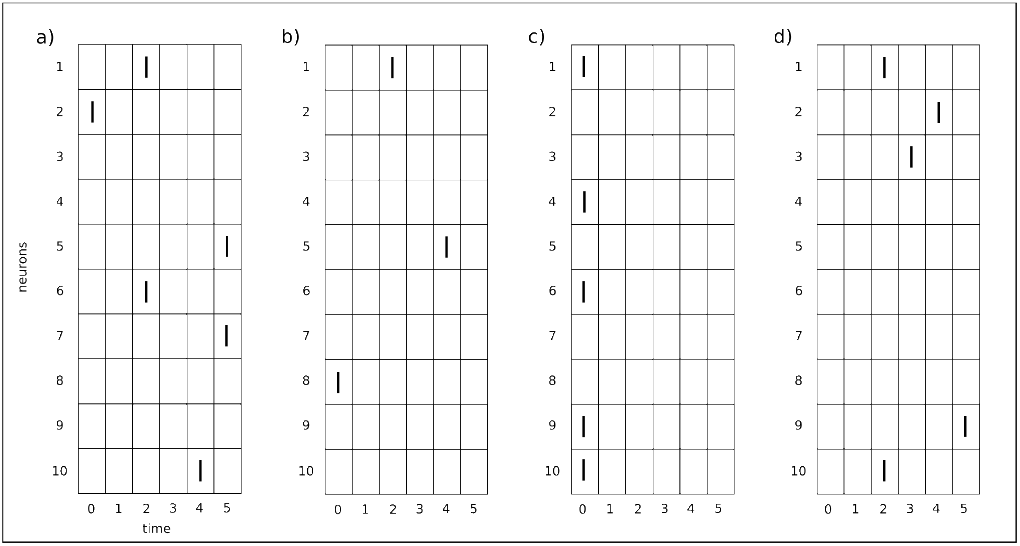
Schematic representation of spike patterns and STPs. In each panel, time indices (𝒲 *=* {0,…, 5}) and neuron indices (𝒩 *=* {1,…, 10}) are indicated by x- and y-axis, respectively. (**a**) *N* (*P* ) *=* {1, 2, 5, 6, 7, 10}, *T* (*P* ) *=* {0, 2, 4, 5}, size |*P*| *=* 6, duration *d =* 5. This is a spike pattern but not an STP because |*N* (*P* )|≠ |*P*|. (**b**) *N* (*P* ) *=* {1, 5, 8}, *T* (*P* ) *=* {0, 2, 4}, size |*P*| *=* 3, duration *d =* 4. This is an STP. (**c**) *N* (*P* ) *=* {1, 4, 6, 9, 10}, *T* (*P* ) *=* {0}, |*P*| *=* 5, *d =* 0. This is also an STP. (**d**) *N* (*P* ) *=* {1, 2, 3, 9, 10}, *T* (*P* ) *=* {2, 3, 4, 5}, |*P*| *=* 5, *d =* 3. This is not an STP because min *T* (*P* ) *=* 2≠ 0.

Mathematically, patterns of size 2 can be considered as a borderline case, and all our definitions and theorems apply to them also. Intuitively, however, they would hardly be called patterns, and in neuroscience, they have been analyzed extensively in terms of pairwise-correlations between spike trains. Here, we are interested in higher than second-order statistics, so we will only treat patterns of size larger than 2.

### Definition 3

(Pattern repetition event) Let 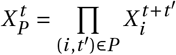 ; it indicates the occurrence of an STP *P* ∈ 𝒫 at time 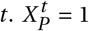 exactly when the variables 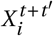 are all 1, i.e., all the neurons involved in the STP *P* emit spikes at the respective times *t + t* ^*′*^. For a given set 𝒮 ⊆ 𝒯 with |𝒮 | *>* 1, we define 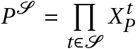 which indicates a **Pattern Repetition (PR)**. The set 𝒮 ⊆ 𝒯 is the set of occurrence times of the pattern *P* . We call the event [*P*^𝒮^ *=* 1] a **Pattern Repetition Event (PRE)**. We also define the set *π* of all PRs by

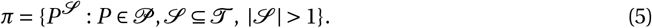

The quantity *r =* |𝒮 | is called the **repetition** of the PR or the number of occurrences of the pattern *P* . In addition we define the random variable

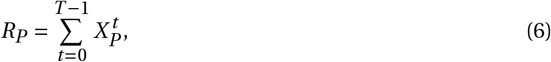

which is called the **repetition of** *P* .

Note that *R*_*P*_ (*x*) is the number of occurrences of the pattern *P* in the spike train *x* ∈ Ξ. The occurrence time of a pattern *P* is the time of its first spike. The concepts of a PRE is illustrated in Fig. 2.

**Fig. 2:**
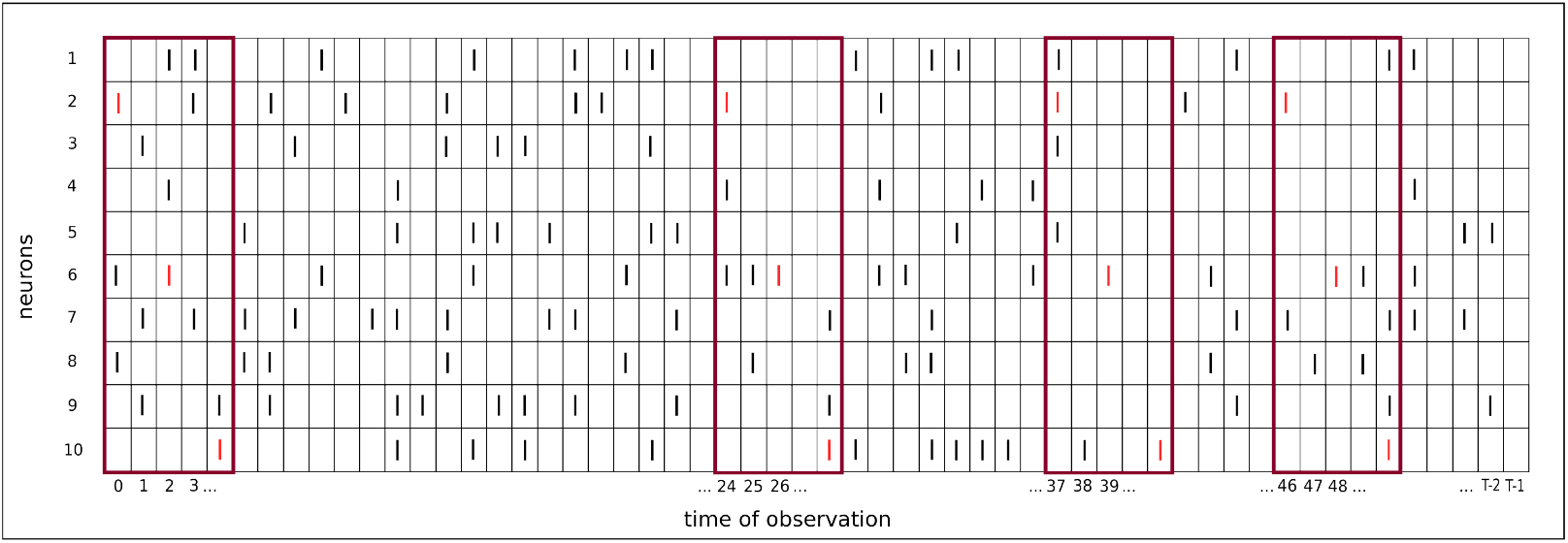
Schematic representation of a PRE (size 3, duration 4, repetition 4, occurrence times 𝒮 *=* {0, 24, 37, 46}.) in a MUST with *N =* 10 units and *T* time bins. Spikes in the repeating pattern are shown in red, and the windows containing the pattern are indicated by red rectangles.

It is important to realize that, for the occurrence of a spike pattern at a particular time, only the occurring spikes matter, not the non-occurring ones, i.e., only the ones in the MUST, not the zeros. This means, for example, that between the first and the second spike in the pattern, there may be additional spikes occurring in the neurons of the pattern. This focus on the occurrence of spikes vs. the non-occurrence has to do with the sparsity of neural spiking and the much higher information contained in the ones versus the zeros in a MUST, and it often goes without saying in neuroscience.

Sometimes it may be reasonable to ask for some separation in time between successive occurrences of an extended STP.

### Definition 4

(Minimal separation) Given a set 𝒮 ⊆ 𝒯, the **(minimal) separation** of 𝒮 is the quantity

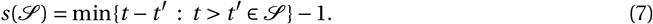

### Definition 5

(Novelty) Given a pattern repetition *P* ^𝒮^, the **novelty** of *P* ^𝒮^, or of the event [*P* ^𝒮^ *=* 1] is defined as

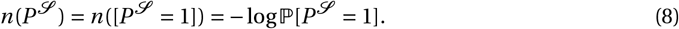

In addition, we define the (average) **pattern novelty** of the repeating pattern *P* as

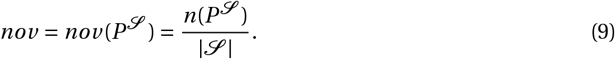

We also define 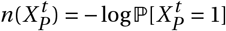 and the random variables

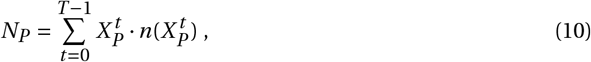

called the **repetition novelty** of *P*, and

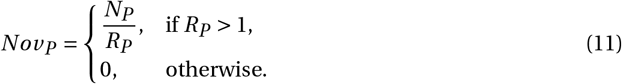

called the **pattern novelty** of *P* .

The definition of novelty as a random variable given here follows the definitions given in Palm (2012) because *N*_*P*_ is the novelty of all possible PREs of a particular pattern P.

Having defined STPs and PRE, we now want to define a quality measure for them. The general idea is to base such a measure on some intuitive criteria for how interesting, surprising, or conspicuous an STP, or rather a PRE, may be. The minimal structure that is needed for this is the ability to compare STPs with respect to these criteria, which is mathematically represented by a ‘≥’ relation. We don’t even assume that any two STPs can be compared, nor do we exclude the possibility that two different STPs may obey the ≥ relation in both directions. The weakest possible relation of this kind is a so-called *preorder*, i.e., a reflexive and transitive relation. In terms of this preorder, we define the ‘cone’ based on a PR *P*^*S*^ as the set of all STPs *P*^*′*^ that occur at times *S*^*′*^ such that *P*^*′S*^*′* ≥ *P*^*S*^ . We try to estimate the number of STPs *P*^*′*^ that have a PR in this cone. This is the subject of the next two definitions of ubiquity and quality of a PR or PRE.

In our previous practice (Stella et al. 2019), we had a slightly simpler problem, as we had some reasonable numerical quality criteria for STPs or PRs, but we did not know how to combine them into a single criterion. The quality measure that we introduce here provides a principled way of combining such criteria. Since it is based on the number of PRs *P*^*S*^ of a certain type, it can also serve as a statistical evaluation of a PRE in terms of the likelihood of observing ‘such an event’, i.e., of a PR in the ‘cone’ based on *P*^*S*^ (see below).

Here we use the term ‘likelihood’ in a non-technical sense, for example, when we tend to assume a ‘meaning of’ or ‘reason for’ something we observe, when it is very unlikely. This intuitive idea has been formalized in several ways in statistics, mostly in terms of probability or conditional probability. In the case of patterns or pattern repetitions, it is particularly difficult to obtain a good estimate for likelihood, because each particular pattern or pattern repetition usually has a very low likelihood, but there are very many potential patterns that would also appear as conspicuous or surprising, and one should actually consider the likelihood of observing any of them. This problem is formalized in Palm (2012), resulting in the distinction between novelty and surprise. Here, we use the ubiquity of a pattern to formalize the more intuitive idea of likelihood. This means that after observing a particular PRE [*P*^*S*^ *=*1], the ‘likelihood’ of observing ‘such an event’ is formalized as the expected number of STPs in the ‘cone’ based on *P*^*S*^, i.e., the expected number of patterns that are at least as surprising as *P*^*S*^ .

### Definition 6

(Cone based on a PR) Consider a given preorder ≤ on the set *π* of all PRs. For a pattern repetition *P*^𝒮^ and a particular MUST x, we define the **pattern cone based on the PR** *P*^𝒮^ as

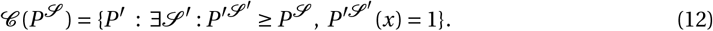

We define the random variable *U* (*P*^𝒮^ ) *=* |*C* (*P*^𝒮^ )| as the number of patterns in this cone. *U* (*P*^𝒮^ ) is a random variable because it depends on a particular spike train *x* ∈ Ξ. The expectation of this random variable is the **Ubiquity** of the PR in which the cone is based.

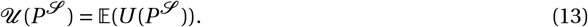

In addition, the **Quality** of a PR is defined as

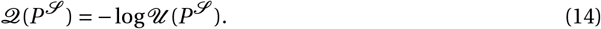

In the next section, we will show how the ubiquity and the quality of a PR can indeed be calculated for one specific preorder, at least for stationary processes. In fact, it can be calculated more or less in the same way for many preorders as long as one is able to enumerate the patterns in the pattern cone. In Section 4, we will discuss the choice of different pre-orders.

The quality of a PR defined here can be used to compare repeating patterns across different signatures, e.g., sizes, repetitions, and also novelties. It can also be used for summary statistics of the overall MUST in order to argue for the existence of repeating patterns beyond chance. In this respect, one could argue for a surprisingly large number of PRs with a large number of occurrences, for example, but without particular regard to their quality (as in the early papers on repeating spike patterns, e.g. Villa and Abeles (1990); Prut et al. (1998)) or, at the other extreme, for the existence of just one or a few patterns of surprisingly high quality (Def. 7). This is the approach we prefer here, because it can also be used to evaluate individual patterns. In a way, one can also combine these two aspects in terms of an adjustable definition of high-quality patterns (see Def. 8).

### Definition 7

(Maximum quality and top-k quality) Given a MUST process, we define the random variable

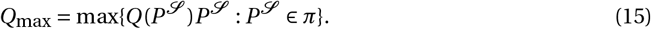

*Q*_max_ is called the **maximum quality** of the MUST.

We also define the **top-k quality** in terms of the quality of the k occurring PREs *PR*_1_,…, *PR*_*k*_ with the highest quality. The random variables *Q*_*i*_ that are the top k qualities, are the rank ordered variables of the set ℳ *=* {*Q*(*P*^𝒮^ )*P*^𝒮^ : *P*^𝒮^ ∈ *π*} and can be defined recursively as follows: ℳ_1_ :*=* ℳ, *Q*_1_ :*=* max ℳ_1_, ℳ_2_ :*=* ℳ_1_ \ *Q*_1_, *Q*_2_ :*=* max ℳ_2_, and so on.

*Q*_*k*_ is called the **top-k quality**.

Using the maximum quality *Q*_*max*_ of a MUST, for the statistics, is analogous to the idea explained in Palm (2012), to use the ‘surprise’ defined there. In view of this analogy, we could call *S*(*x*) *=* − log(*p*[*Q*_*max*_ ≥ *Q*_*max*_ (*x*)]) the pattern repetition surprise of *x*. A different idea is based on the definition of high-quality PRs by means of a quality threshold instead of just considering the top k PRs.

### Definition 8

(High-quality PRs) Given a MUST process, a quality threshold *q*_−_ ∈ **ℝ**, and a repetition threshold *r* ∈ **ℕ**, we define the set of **high-quality PRs** as

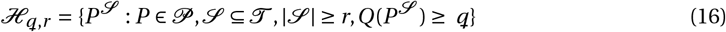

Then we define the random variables

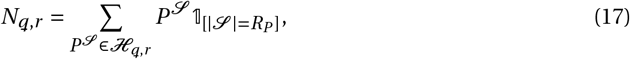

and

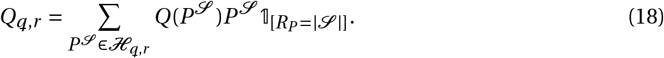

*N*_*q,r*_ is the number of high quality PRs beyond a given quality threshold *q* and *Q*_*q,r*_ is their total summed quality. The quality measure makes it possible to adjust this threshold between the two extremes of considering large numbers of PRs and one PR of highest quality. In many experimental paradigms, in particular when the same condition is repeated several times, it also makes sense to introduce a threshold *r* on the repetition.

The distribution of the variables defined here can then be used for statistical testing of a MUST. However, it is practically impossible to determine this distribution mathematically. Therefore, one will have to resort to extensive sampling. We will come back to this issue in Section 5.

## 3 Computing quality

### 3.1 Mathematical estimation of ubiquity

We first define the preorder ≥ between PRs. The criteria that we use for this have essentially all been defined in the last section. They are the *size z*, the *duration d*, the *repetition r*, and the *pattern novelty nov* (or *novelty n*); one could also consider the *separation s*. In principle, these parameters can also be considered as properties of repeating STPs, but then we have to distinguish between **structural** parameters describing the pattern itself, namely *z* and *d*, and **probabilistic** parameters describing its occurrences, which are random variables, namely *R*_*P*_ (for *r* ) and *Nov*_*P*_ (for *nov* ; or *N*_*P*_ for *n*). Previously, *z, d*, and *r* were used in SPADE (Quaglio et al. 2017), where *z* and *r* are inherited from frequent itemset mining (Borgelt and Picado-Muiño 2013b,Borgelt and Picado-Muiño 2013a; Borgelt 2012), and the novelty *n* was introduced in Palm (2012). All these parameters clearly affect the likelihood of pattern repetitions. Larger size *z*, larger repetition *r*, and of course larger pattern novelty *nov* will intuitively (and also mathematically as we will see) decrease the likelihood, whereas larger duration *d* will increase the likelihood for *z >* 2, because then there are more possibilities to put additional spikes between the first and the last spike of a pattern.

In the general non-stationary setting, it may sometimes be useful to define an additional criterion which can be viewed as a simpler substitute for the pattern novelty *nov* . We suggest a substitute that is based on the average firing rates of the neurons in a pattern and, therefore, is not a random variable. We call it the *simple novelty*.

#### Definition 9

(Simple Novelty) The **Simple Novelty** of a pattern *P* is

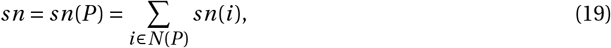

where 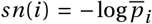 with 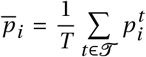 being the average probability of firing of neuron *i* . By definition, it is a function of the overall firing rate of the neuron.

Furthermore, *sn*(*P* ) depends only on the neurons *N* (*P* ) in *P*, not on the time delays that define the pattern *P* .

Now we define a preorder ≥ between PRs in terms of some of these parameters, which we use for this paper. We collect all the numerical quality criteria that are needed to define ≥ into a vector, which we call the **signature** *σ* of a PR *P*^𝒮^ . For the following discussion, we take just 3 parameters, omitting the size *z*:

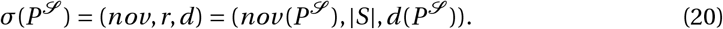

#### Definition 10

(Preorder) Given two signatures *σ =* (*nov, r, d* ) and *σ*^*′*^ *=* (*nov*^*′*^, *r* ^*′*^, *d*^*′*^) we say that *σ*^*′*^ ≥ *σ* when

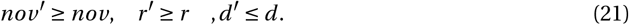

Based on this we can define a **preorder** ≥ also on PRs by simply defining 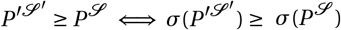

This means that in the simple situation where a preorder on PRs can be defined in terms of numerical criteria collected in a signature, this preorder can already be defined on the space of signatures, where it even is a partial order (i.e. *σ* ≤ *σ*^*′*^ and *σ*^*′*^ ≤ *σ* implies *σ = σ*^*′*^). So, the cone, the ubiquity, and the quality can also be defined for signatures. In addition, we can easily define slightly different pre-orders based on different combinations of our quality criteria, by simply listing them in the signature and defining an order on each quality criterion. For example, we can replace *nov* by *sn*, or by *z*, or we could add *z* to the signature, using *σ =* (*sn, r, d* ) or (*z, r, d* ) or (*nov, z, r, d* ), resulting in different versions of ubiquity and quality.

#### Definition 11

(Cone based on a signature) Given a signature *σ*, the **pattern cone based on the signature** *σ* is defined as follows

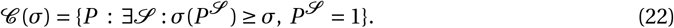

Let *U*_*σ*_ *=* |𝒞 (*σ*)| be the number of patterns in this cone. *U*_*σ*_ is a random variable because it depends on a particular spike train *x* ∈ Ξ. The expectation of this random variable is the **Ubiquity** of *σ* and also of any PR with this signature.

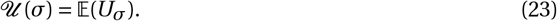

Again, the **Quality** of *σ* is

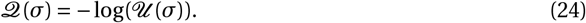

Fig. 3 illustrates the concept of the cone based on a signature.

**Fig. 3:**
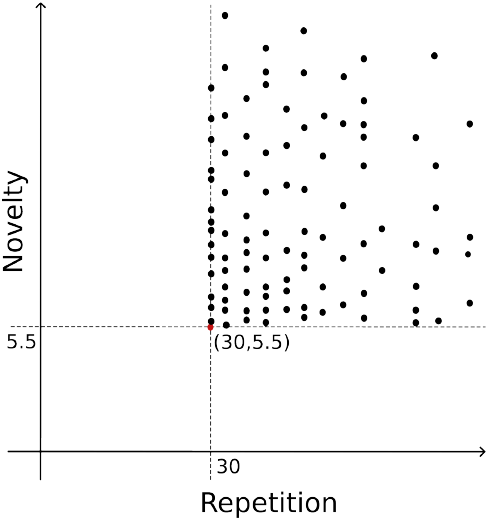
Schematic representation of a cone obtained from a MUST, defined by a signature *σ =* (*nov, r, d* ) *=* (5.5, 30, *τ*), or equivalently a PR with the same signature. The pattern duration *d* is fixed to *τ*, leaving pattern novelty (*nov* ) and pattern repetition (*r* ) as the displayed dimensions. The dashed lines indicate the cone boundaries, and the red point marks the base of the cone at (*r, nov* ) *=* (30, 5.5). Black dots represent PRs (in the MUST) contained in the cone, satisfying *nov* ≥ 5.5, *r* ≥ 30, and *d* ≤ *τ*.

Based on this definition, we can derive the formulas for the ubiquity and the quality of a PR, or even an STP, with a given signature, when the stochastic process ***X*** is stationary. We start by putting together the simplifications provided by assuming stationarity.

#### Proposition 1

*If the process X is stationary, i*.*e. if* 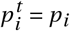 *for all t* ∈ 𝒯, *then for any STP P we have*

1. 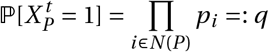*and, for a PRE in which P occurs r times, we have* 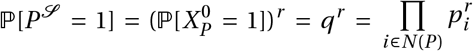.
2. 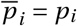 *for all i, nov = sn and N*_*P*_ *= R*_*P*_ · *sn*.
3. *R*_*P*_ *is binomially distributed, i*.*e*., 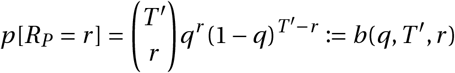, *where T* ^*′*^ *is defined in Prop. 2*.

*T* ^*′*^ represents the number of time-points at which a pattern can occur. It is typically slightly smaller than *T*, depending on the pattern duration d. In general, T’ is equal to *T* −*d* .

Now we make some simple combinatorial observations.

#### Proposition 2

*It holds that*

1. *The number of occurrence time sets* 𝒮 *of PRs with minimal separation s, repetition r, and duration d is*

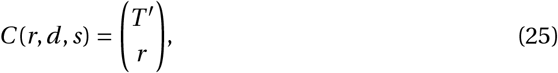

*where T* ^*′*^ *= T* −*d* − *s*(*r* − 1).
2. *The number of spike patterns of size z and duration* ≤ *d, starting at time 0, is*

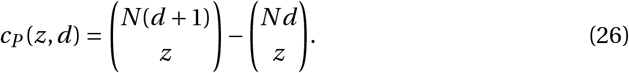
3. *The number of STPs of size z and duration* ≤ *d is*

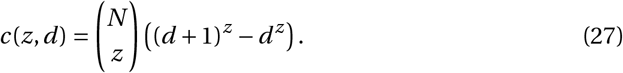

*Proof* We prove the three statements in turn.

1. We can map subsets *R* ⊆ {0,…, *T* − (*r* − 1)*s*} of size *r* into subsets *S* ⊆ {0,…, *T* } with separation s in a 1-to-1 fashion: For an ordered sequence 0 ≤ *t*_1_ *< t*_2_ *<* · · · *< t*_*r*_ ≤ *T* − (*r* − 1)*s* we define *f* (*t*_*i*_ ) *= t*_*i*_ *+* (*i* − 1)*s* to get 0 ≤ *f* (*t*_1_) *< f* (*t*_2_) *<* · · · *< f* (*t*_*r*_ ) ≤ *T* and { *f* (*t*_1_), …, *f* (*t*_*z*_ )} have separation s. Therefore, the number of subsets of size r with separation s(from {0,)…, *T* } equals the number of subsets of size r from {0,…, *T* − (*r* − 1)*s*} which is 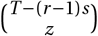.
2. A spike pattern of duration *d* is an arbitrary subset of the set {1,…, *N* } *×* {0,)…, *d* }, which has size *N* (*d +* 1). Thus, the number of spike patterns of size z is 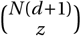. If we want to avoid multiple counting of a pattern, we require that spike patterns (or rather spike pattern templates) start at time 0. This means that we exclude all patterns that start at time 1 or later, i.e., which are ⊆ {1,…, *N* } *×* {1,…, *d* }. Their number is 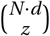, which has to be subtracted.
3. For STPs, we require in addition that each neuron appears only once in a pattern, i.e., we could choose first a subset of z neurons from {1,…, *N* } and then choose for each neuron a time *t* ∈ {0,…, *d* }. This gives 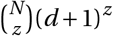 possibilities. Again, we exclude patterns which start at time 1 or later, i.e., where *t* ∈ {1,…, *d* } for each *t* ∈ *T* (*P* ); their number has to be subtracted.

Now we can compute the ubiquity of a signature *σ =* (*sn, d, r* ), which is greatly simplified by taking *nov = sn*, such that *r = R*_*P*_ remains the only random variable in the signature, and we can separate the counting of patterns *P* from the counting of subsets 𝒮 of 𝒯 with |𝒮 | ≥ *r* . And the latter is obtained from the binomial distribution.

To simplify the writing in the next theorem, we briefly introduce an additional notation, namely

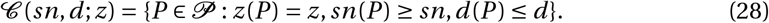

#### Theorem 3

*Assuming stationarity of* ***X***, *the Ubiquity of a PR P*^𝒮^ *or, equivalently, the Ubiquity of the signature σ = σ*(*P*^𝒮^ ), *is*

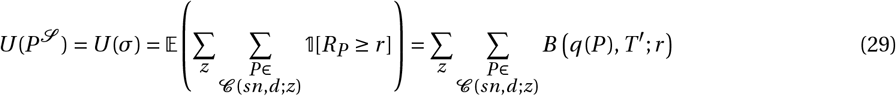

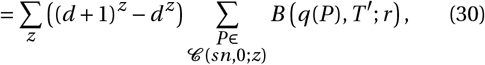

*where B* (*q*(*P* ), *T* ^*′*^; *r* ) *is the Cumulative Binomial distribution*

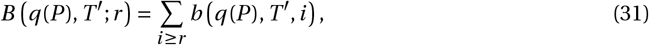

*and*

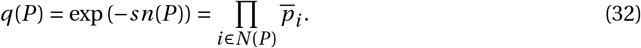

Computationally and numerically, it is still very complex to calculate this formula because it is a very large sum of extremely small terms. For this reason, it is useful to derive a simpler approximation for it. Here we provide a lower and upper bound for the binomial distribution and thereby for the ubiquity. Then we can calculate the ubiquity of a PR or of its signature for the stationary case.

#### Proposition 4

*It holds that*

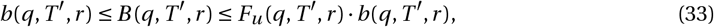

*where the first inequality holds by definition and the second requires that r* ≥ *λ* :*= T* ^*′*^*q. Here*

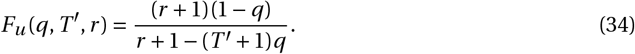

*Proof* This is a well-known estimate of the tail of the binomial distribution by a geometric series:

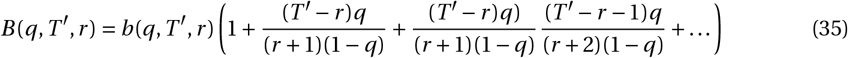

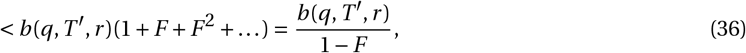

for 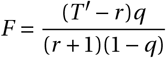.

#### Proposition 5

*Conversely, if* 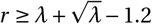, *it holds that*

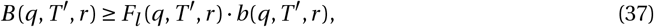

*where*

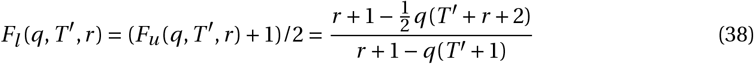

*We also observe that F*_*u*_ /*F*_*l*_ *<* 2 *for r* ≥ *λ*.

*Proof* The proof is given in Appendix A.

To our knowledge, this lower bound to the binomial tail is new. It ensures that the difference between the upper and lower bound of quality stays below log(2), and requires only a weak condition on *r* −*λ*.

An efficient computation of the formula for the ubiquity is provided by putting these estimates (Propositions 4 and 5) together in the following theorem, where we can restrict the necessary summations by assuming that the neurons are indexed in the order of increasing firing probability, and by introducing an additional term: given an increasing sequence of neuron indices *i*_1_ *< i*_2_ *<* · · · *< i* _*j*_, ( *j < z*) we define *sn*_*max*_ ( *j* ) as the maximal value of *sn* that can be achieved after these *j* indices have been fixed, i.e. 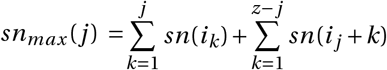. Then we can restrict the summation over *i* _*j*_ by the condition that *sn*_*max*_ *(j)* _*≥*_ *sn*. For this purpose we define 𞟙_*j*_ :*=* 𝟙[*sn*_*max*_( *j* ) ≥ *sn*].

#### Theorem 6

*Assuming stationarity, for a given signature σ =* (*sn, r, d* ) *with* 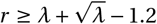 *and λ = T* ^*′*^ · *e*^−*sn*^ *we obtain*

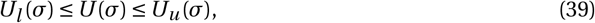

*where*

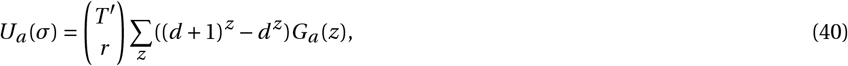

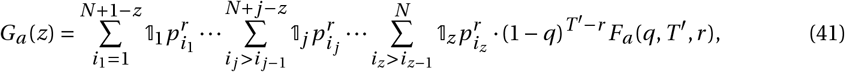

*for a* ∈ {*l, u*} *and* 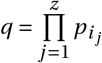.

*G*_*a*_ (*z*) *decreases fast with z; see Appendix B for details*.

*U*_*l*_ and *U*_*u*_ are the lower and upper bounds of the ubiquity. From Prop. 5 we know that *U*_*u*_ /*U*_*l*_ ≤ 2. The corresponding bounds for the quality are *Q*_*l*_ *=* − log*U*_*u*_ and *Q*_*u*_ *=* − log*U*_*l*_ . Thm. 6 concludes our mathematical efforts to compute the quality of a PR.

Now one could be tempted to use the quality and ubiquity of a repeating pattern *P* as defined and calculated here, directly for statistical significance testing, in particular when 𝒰 (*P* ) is small, for example, less than 0.05. Indeed, the probability *p =* ℙ [𝒰 (*P* )≠ 0] ≤ 𝒰 (*P* ). However, this is misleading because *p* cannot be considered as a significant probability, since it depends on one particular pattern (or on the pattern cone based on it), not on all possible cones or the whole MUST. Intuitively, the ubiquity can be used to estimate the probability (under the null hypothesis) of generating a pattern in the cone based on *p*. But there are many more such cones with about equally low ubiquity, which would result in the same or even higher quality. So the question to be asked is: *what is the probability of generating a pattern with a ubiquity U*^*′*^ ≤ *U or with a quality Q*^*′*^ ≥ *Q?* In Palm (2012) the negative logarithm of this probability is called the **surprise**, here of the pattern *P* .

What we are suggesting here is often done in the statistical analysis of events that happen not just once, but many times or in many places in an extended observation, resulting in many values of a potential statistical test variable (like our quality). Instead of evaluating one particular event, one evaluates the whole observation in terms of the total collection of events that happened, but uses a measure that can also be applied to just one event. Then this measure is also used (or kind of abused) to test one particular event In the case of STP quality, we are interested in high values of quality (rather than large numbers of patterns) and so it appears natural to consider the maximal quality achieved in a MUST as our statistical test (see Def. 7). This means,we compare the quality of one particular experimental STP with the maximal quality across all STPs that can be achieved in a simulated MUST.

In addition, the assumption of stationarity that is needed for our calculations is usually too unrealistic, even for a null hypothesis. Therefore we now have to consider the estimation of ubiquity and quality from simulated or surrogate data which are nonstationary.

### 3.2 Simulation-based estimation of ubiquity

To estimate the ubiquity of various PRs based on simulations, one has to generate a large number of realizations of the MUST process (Def. 1). For each MUST realization *x*, one has to find all potentially interesting PRs by, for example, the frequent itemset mining algorithm (Borgelt 2012) and determine their signatures. These signatures are counted and accumulated over all realizations *x* in a signature histogram (here it is a 4-dimensional histogram spanned by size *z*, repetition *r*, duration *d*, and simple novelty *sn* or novelty *nov* ). Note that, since the novelty measures take continuous values, to construct the histogram one has to set discrete bins over the novelty values with a reasonably small bin size; here we take 0.02. For each bin in the histogram, which corresponds to a particular signature *σ*, the counts are summed over all bins corresponding to any signature *σ*^*′*^ satisfying *σ*^*′*^ ≥ *σ*. Dividing the resulting count by the number of realizations of the MUST yields an estimate of the ubiquity 𝒰 (*σ*) of the signature *σ*, and the logarithm of this ubiquity value is the estimate of the quality 𝒬 (*σ*). Here, we take the logarithm to the base 10, which simplifies the reading of the underlying ubiquity values. With this calculation we obtain a 4-dimensional matrix (spanned by the same quality criteria as for the signature histogram) of quality values, which we call a quality spectrum (see Fig. 12 for an example).

In order to achieve estimation of quality up to 4, one needs to accumulate signatures over 10,000 realizations of a MUST. Even with this large number of realizations, estimation of large quality values, say greater than 3, becomes quite inexact due to the small number of corresponding patterns occurring in the 10,000 realizations (quality of 3 corresponds to ubiquity of 10^−3^, i.e., the relevant PRs found only 10 times in 10,000 MUST realizations).

For generating realizations of a MUST by simulation, we need the defining parameters: the recording time *T*, the number *N* of neurons, and the firing probabilities 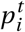, which are derived from experimental spike train data as explained in Section 3.3. Based on those parameters, artificial spike data are generated by randomly drawing a spike with probability 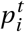 for each bin, which results in a MUST process, i.e., a collection of discretized non-homogeneous Poisson processes.

Alternatively, one could consider surrogates of experimental spike trains (Stella et al. 2022) as realizations of a MUST. Here the firing probabilities 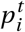 do not need to be estimated from the data. Still, one needs an enormous computational effort to find all interesting spike patterns in each of the surrogates. So the generation of a quality spectrum is very costly.

### 3.3 Estimating parameters from experimental data

To compute the quality according to the formula in Thm. 6, we have to specify the parameters characterizing the spike train data: the recording time *T*, the number *N* of neurons, and a vector *p*_*i*_ of *N* firing probabilities. Estimating the quality from simulations even requires all *T × N* bin-firing probabilities 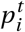. The values of these parameters should be chosen based on the features of experimental data. In the present study, we use data that were recorded in a previous experiment (Riehle et al. 2013), which is briefly described here.

The spike-train data were recorded by a chronically implanted 96-electrode Utah array (Blackrock Microsystems) from the motor cortex of a female monkey (monkey L) performing an instructed-delay reach-to-grasp task (Riehle et al. 2013). The monkey was trained to reach to an object to grip and hold it with different grip types (side or precision grip) and different holding forces (high or low force). We concentrate here on the precision grip (PG) and high force (HF) conditions only.

Fig. 4 (top) shows the raster plot of the spike trains of 88 simultaneously recorded neurons in an example PGHF trial, with the times of the behavioral events. Fig. 4 (bottom) displays the firing probability (per 5 ms bin) of each neuron estimated from the spike train in the top plot. An experimental trial contains different behavioral phases that directly influence the firing probabilities of the recorded neurons. To derive the data-based parameters for the evaluation of the quality according to the formulas in Thm. 6 that assume stationarity, we focus on the period with relatively stationary firing rates, i.e., the ‘wait’ period (500 ms long), when the monkey just waited for the first cue around the onset of waiting signal (WS-ON). Later, we will further discuss the validity of the stationarity assumption by comparing the quality estimation from stationary and non-stationary spike train data. For that purpose, we will also focus on the period with non-stationary firing rates, i.e., the ‘movement’ period around the movement start (‘switch release’, SR). These two periods are indicated by the colored shades (wait: orange; movement: purple) in the figure.

**Fig. 4:**
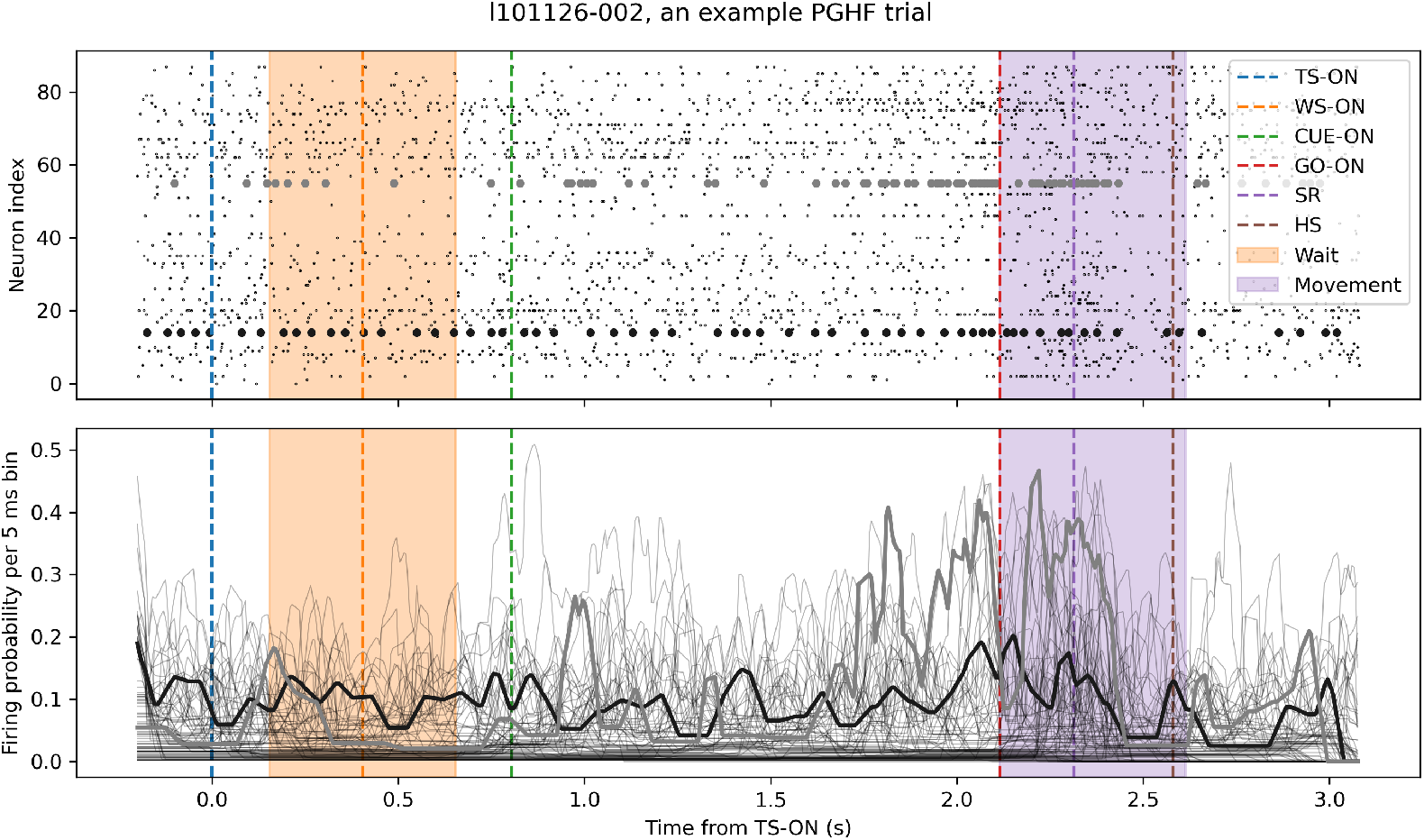
Example of experimental data. Top: Raster display of the spiking activity of all 88 simultaneously recorded neurons during one trial (out of 36) of trial type PGHF in session l101126-002. Each dot shows the occurrence of a spike for each of the neurons (y-axis). The dashed vertical lines mark the points in time when the trial starts (TS-ON, blue), the waiting signal occurred (WS-ON, orange), and when the cue started (CUE-ON, green). GO-ON (red) indicated that the monkey should start to move the arm to grasp the object, and at SR (switch release) he actually started to move. At HS, the monkey started to hold the object. For the analysis performed here, we selected two periods of 500 ms duration from each trial, one around WS-ON (+/- 250 ms) (‘Wait’) and during movement starting around SR (-200 ms, +300 ms) (‘Movement’). The spiking activities of 2 neurons are illustrated with thicker dots to show more clearly the changes in the firing rates. Bottom: Instantaneous firing probabilities for each of the neurons in the same trial as the data shown above. For illustration of the potential differences in the time-dependent firing rates, two firing rate profiles are highlighted in thicker lines.

For the evaluation of the quality under stationarity assumptions, the firing probabilities *p*_*i*_ were estimated from the average firing rate per neuron per period (in wait or in movement, separately), assuming stationarity. In the more general non-stationary case, we need the firing probability 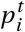 for every neuron *i* and every time-bin *t* . These ‘instantaneous’ firing probabilities are calculated in the following way. First, the inter-spike intervals (ISI) are computed between any two successive spikes of a neuron. The inverse of that (1/ISI) is considered as an estimate of the instantaneous firing rate (Koštál and Kováčová 2025) during the time period of that ISI. This procedure yields a continuous time series of instantaneous firing rate estimates, with step-wise changes at the timings of the spikes. This time resolved estimate is then binned in time with a bin width of 5 ms, and smoothed by taking the moving average of neighboring 9 bins. The firing probabilities shown in Fig. 4 (bottom) were obtained in this way.

## 4 Basic Properties of Quality and Computational Simplifications

### 4.1 Dependence of quality on signature parameters

Given the data-based parameters derived from the experimental data, we can illustrate how the quality of a PR depends on the parameters of its signature, by computing the quality according to the formula in Thm. 6 and plotting it as a function of the signature parameters. In Fig. 5 we display the quality as a function of each of the parameters *r, d, sn* for *z =* 3. One can see that the quality depends on each of these parameters as intuitively expected: it increases with *sn* and *r*, and it decreases with *d* . Also, the duration *d* has a smaller effect on quality than the repetition *r*, i.e., changing *d* by 1 has a smaller effect than changing *r* by 1.

**Fig. 5:**
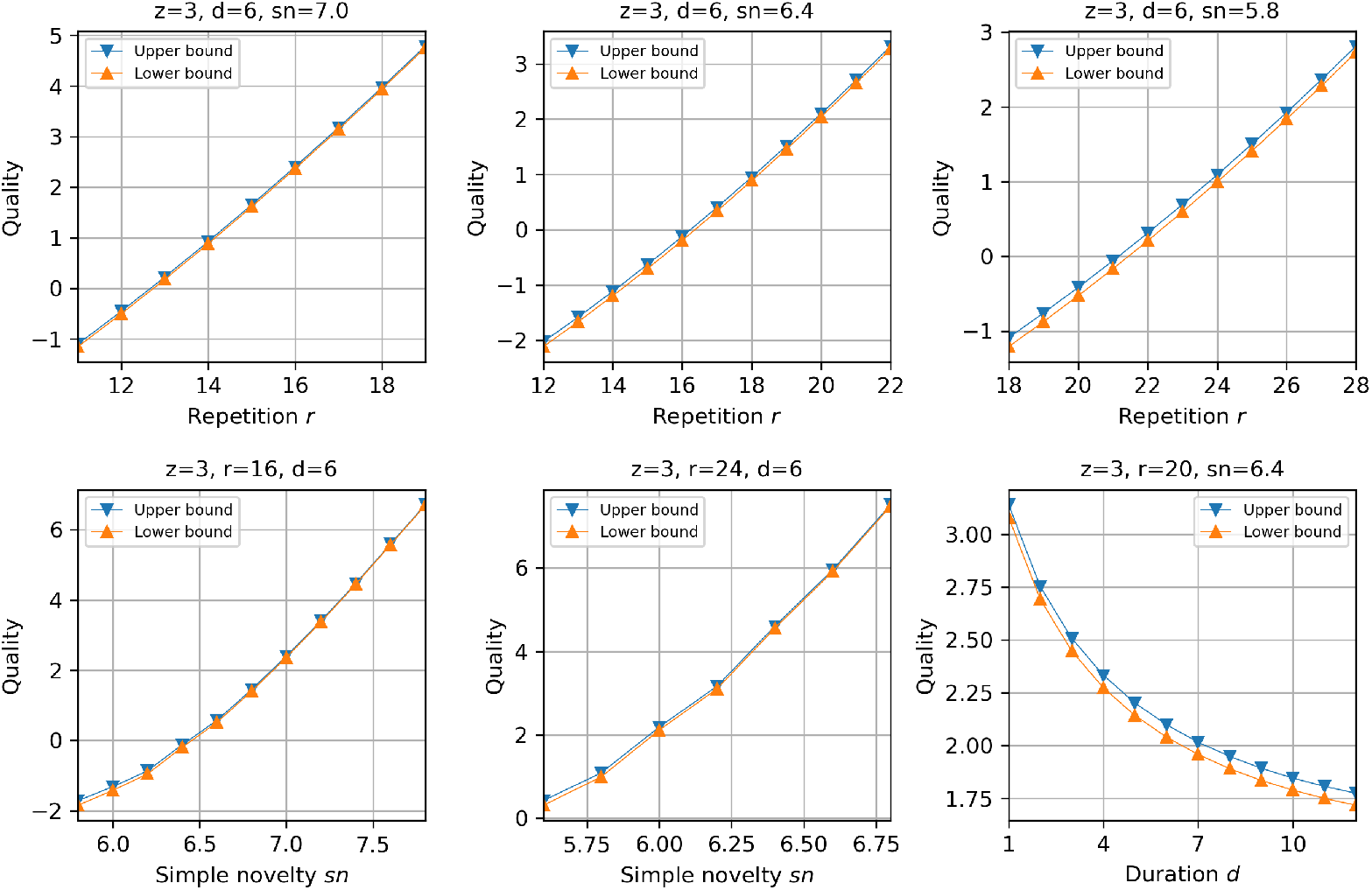
The upper and lower bounds of the quality computed according to Thm. 6. Each panel shows the dependence of the quality bounds on a single signature parameter, with the other signature parameters fixed as indicated above the plot. The other parameters characterizing the underlying spike train (the recording time *T*, the number *N* of neurons, and a vector *p*_*i*_ of *N* firing probabilities) were derived from the ‘wait’ period of PGHF trials in the reach-to-grasp experiment dataset, as described in Section 3.3.

In general, the monotonicity of quality with respect to the parameters in the signature (or to the corresponding pre-order) on which it is based is an essential property that has to be expected for any reasonable quality measure.

Before computing the quality values shown in Fig. 5, we had checked the correctness of our formulas for several simple examples with only 1 or 2 different values for the parameters *p*_*i*_ . To further confirm the correctness in more complex cases, we also simulated a stationary MUST of many spike trains, all with different firing probabilities *p*_*i*_, and estimated ubiquity and quality for various signatures as explained in Section 3.2, to compare the results to the the quality values from the formulas. Concretely, we used the experimental data described in Section 3.3 and generated the simulated data as spike randomization surrogates, i.e., for each neuron, the number of spikes in a period was counted and distributed randomly in time. This procedure generates stationary spike trains.

Fig. 6 (left) shows the comparison of the analytical upper and lower bounds of the stationary quality in the wait period against the corresponding simulation-based estimate, for a set of signatures sampled in ranges of repetition *r*, duration *d*, and simple novelty *sn* (see Appendix E for the complete list of the signatures). Fig. 6 (right) shows the same analysis for the stationary quality in the movement period, which in reality had strong non-stationarity in the firing rates. In both cases, the simulated quality is very close to the upper or lower boundary of the stationary quality obtained from the formulas for most of the signatures. Yet, the diagonal is not always enclosed between the upper and lower bounds (i.e., the simulated quality is within the computed bounds), but we observe a consistent discrepancy towards higher values of the simulated quality versus the computed quality, which increases for higher quality values. This increase and also the increasing variation towards higher quality values may be due to the small numbers of patterns that correspond to quality values larger than 3, as mentioned in Section 3.2. The reason for the small, but consistent discrepancy could perhaps be the extensive use of pseudo-random numbers, for a somewhat untypical purpose. We did not investigate this discrepancy more deeply, because we are heading now in a different direction, namely the much larger effects of non-stationarity.

**Fig. 6:**
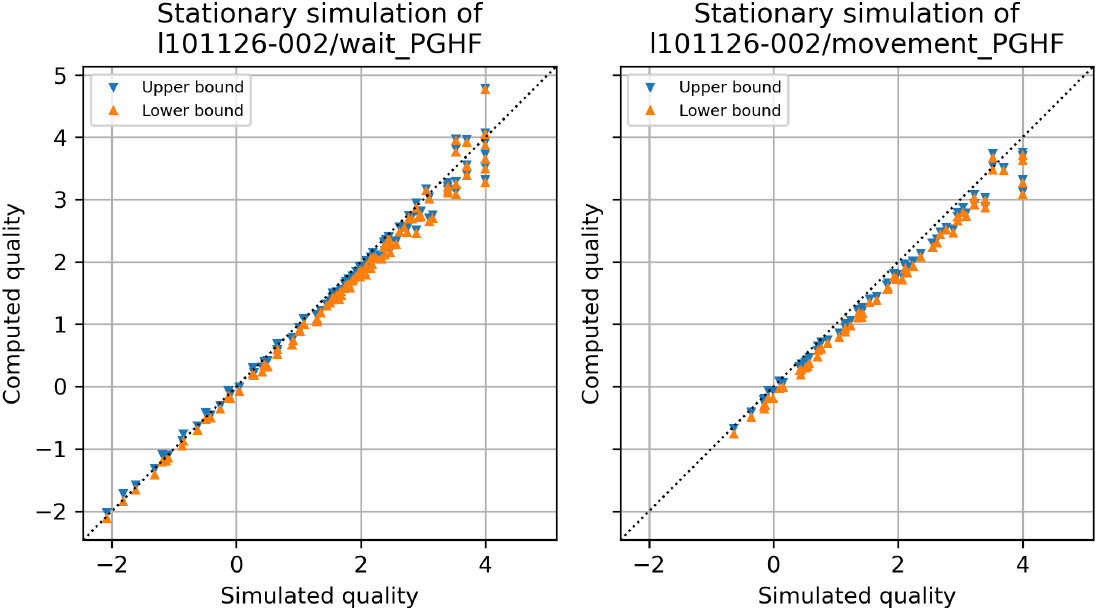
The quality bounds computed according to Thm. 6, assuming stationarity (computed quality), are plotted against the quality estimate based on stationary simulations (simulated quality). Each data point in the plot corresponds to a signature sampled in ranges of repetition *r*, duration *d*, and simple novelty *sn* (see Appendix E for the complete list of the sampled signatures). The data-characterizing parameters for the computation and simulation were derived from the ‘wait’ period (left) or the ‘movement’ period (right). For each period, 10,000 simulations (spike time randomization surrogate) were performed to estimate the simulated quality.

### 4.2 Relating stationary and non-stationary quality

We have been able to calculate the ubiquity and quality of any given PR in terms of its particular signature, under the assumption of independence and stationarity. We can also use this measure for the numerical comparison of repeating patterns.

In classical statistics, it is quite common to use such strong stochastic assumptions as a null-hypothesis of complete randomness because this allows the analytical calculation of the probability distribution of the test-statistics. In modern experimental science, however, this is often not sufficient because one is interested in a subtle difference between the new experimental hypothesis and an already quite elaborated “null-hypothesis”. In this situation, it is usually not possible to calculate the distribution of the test variable, but one has to resort to extensive sampling from the null-hypothesis, which is computationally expensive. In fact, often the ‘null-distribution’ is not even defined mathematically (involving just a few parameters from the data), but operationally by the ‘sampling’ procedure, which typically consists of data manipulations that generate so-called ‘surrogate data’ and rely heavily on the experimental data (Lancaster et al. 2018; Stella et al. 2022).

In our case, in the multiple-unit spike-trains, it is in particular the assumption of stationarity that is problematic. In recordings from a living animal, non-stationarities naturally arise from external stimuli or internal influences from the animal’s body or other brain areas, but also from unwanted changes in the recording conditions. We hypothesize that the patterns we are looking for are due to (synaptic) interactions between the participating neurons, and most likely also to spike inputs from other non-recorded neurons, and we want to exclude the effects by covariation of firing rates due to external stimuli or due to the relation to movements. For this purpose, it makes sense to assume independence at least between the spike trains of different neurons as a null-hypothesis, but not stationarity; this we did in Def. 1. However, one could also consider a more complex ‘data-driven’ null-hypothesis which is based on surrogate data. In both cases, it is not possible to calculate ubiquity or quality directly, but one has to resort to extensive sampling.

In practice, it is often not feasible to employ such a computationally demanding method for every dataset on which one wants to compare patterns according to their quality, or to evaluate the statistical significance or unexpectedness in terms of their quality. In this situation, one may hope to use the ‘stationary quality’ that we computed in the last section, or perhaps an even more easily computable quantity as a kind of ‘proxy’ for quality. It may therefore be interesting to look at the relation between the ‘stationary quality’ and the ‘real’ quality as we defined it in Section 2.

From now on, this calculated ubiquity (and quality), which is based on stationarity, will be called the *stationary ubiquity U*_*s*_, resp. *stationary quality Q*_*s*_ . Since we are assuming stationarity for the calculation of *Q*_*s*_, which implies that *nov = sn*, it seems more natural to use *sn* in the formula that computes quality. In principle, one could also use *nov*, but this would assume that the neurons are always firing with the same (typically higher) frequency that occurs during the repeated patterns.

The ‘real’ quality of non-stationary data needs to be estimated based on simulations as described in Section 3.2. As shown in Fig. 4 (bottom), the firing rates during some phases of the experiment, in particular during the movement period, change a lot. To accurately estimate the ‘real’ quality (Def. 6) of PRs found in such non-stationary data, we need to first estimate the trial-by-trial instantaneous firing rates of the individual neurons and use these estimates to determine the parameters 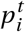 needed for the MUST simulations as described in Section 3.3. Based on these simulations and the quality spectrum generated from them, we can compare (the estimate of) the ‘real’ quality defined in Def. 6 with the computed values of the stationary quality defined in Thm. 6.

Fig. 7 shows the comparison for the wait period (left) and the movement period (right). In the ‘relatively stationary’ example of Fig. 7 (left), we see a reasonable agreement between the two qualities. In the strongly non-stationary dataset of Fig. 7 (right), we observe that the simulated quality Q is clearly smaller than the calculated quality *Q*_*s*_, but there are also some outliers with very high stationary quality and the correlation is not very strong. However, there is still a roughly monotonic relationship between the two quantities.

**Fig. 7:**
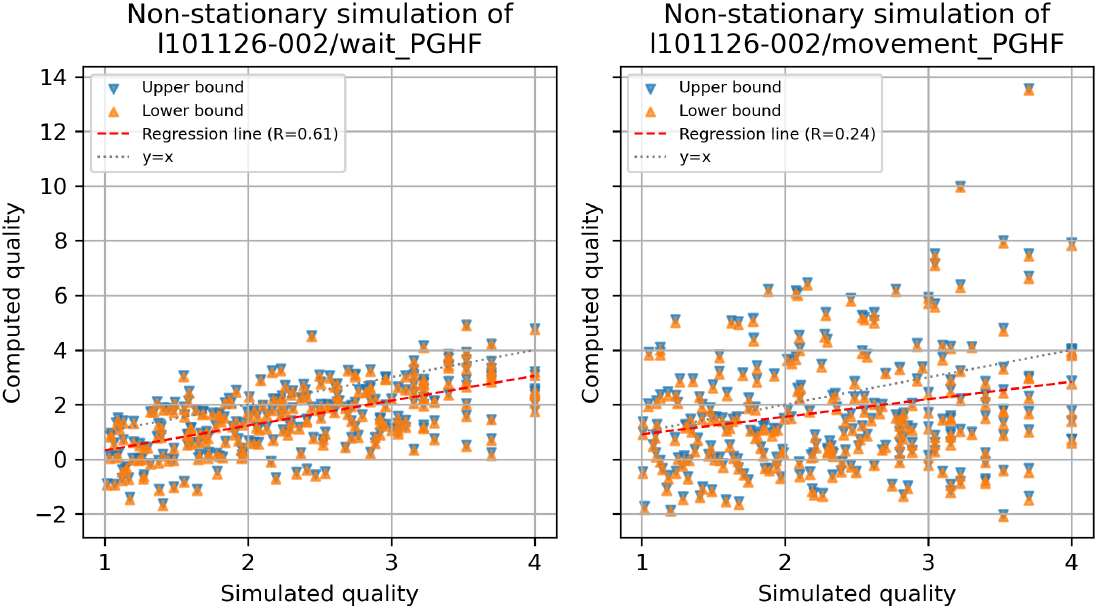
The quality bounds computed according to Thm. 6, assuming stationarity (computed quality), are plotted against the quality estimate based on non-stationary simulations (simulated quality). Each data point in the plot corresponds to a signature sampled in terms of their estimated quality (see Appendix E for how the signatures are sampled). The data-characterizing parameters for the computation and simulation were derived from the ‘wait’ period (left) or the ‘movement’ period (right). For each period, 10,000 simulations of a MUST process with the ISI-based estimation of the (non-stationary) firing probabilities were performed to estimate the simulated quality. The red line indicates the least-square regression line, the correlation coefficient *R* of which is shown in the legend.

When we used the pattern novelty *nov* instead of the simple novelty *sn* in the stationary formula (described in more detail below), we found a less scattered relationship with a higher correlation coefficient (see Fig. 8). Therefore, it may be possible to estimate Q from this version of *Q*_*s*_, or from *Q*_*s*_ itself, at least roughly. And the estimated Q increases monotonically with *Q*_*s*_ . This suggests that for the comparison of PRs, and also for statistical testing, one could use this version of *Q*_*s*_, which can be directly calculated from the pattern signature, as a substitute for the real ‘dynamic’ quality *Q* in the non-stationary surrogate process.

**Fig. 8:**
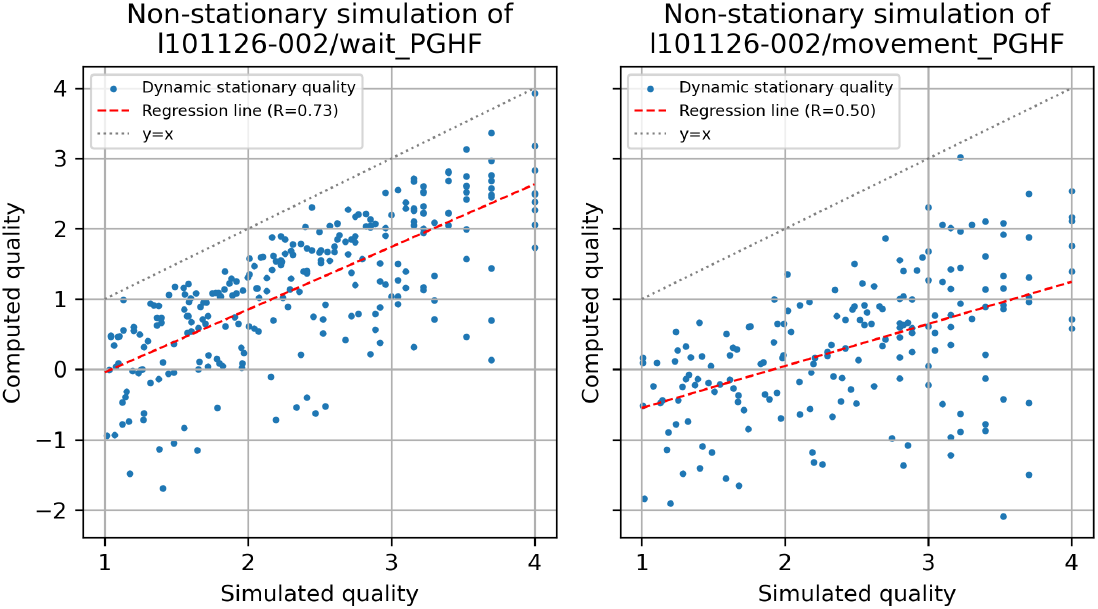
Same plots as in Fig. 7, but with the computed quality using the original novelty *nov* instead of the simple novelty *sn*.

The difference between the use of *sn* and *nov* in the formula for *Q*_*s*_ can be easily explained by the quite consistent difference between the values of *sn* and *nov* themselves (Fig. 9): *sn* is almost always larger than *nov*, which means that the patterns occur in periods where the contributing neurons are firing more intensely than on average. This is the reason why it makes sense to use *nov* in the formula, although its use clearly violates the stationarity assumption on which the formula is built.

**Fig. 9:**
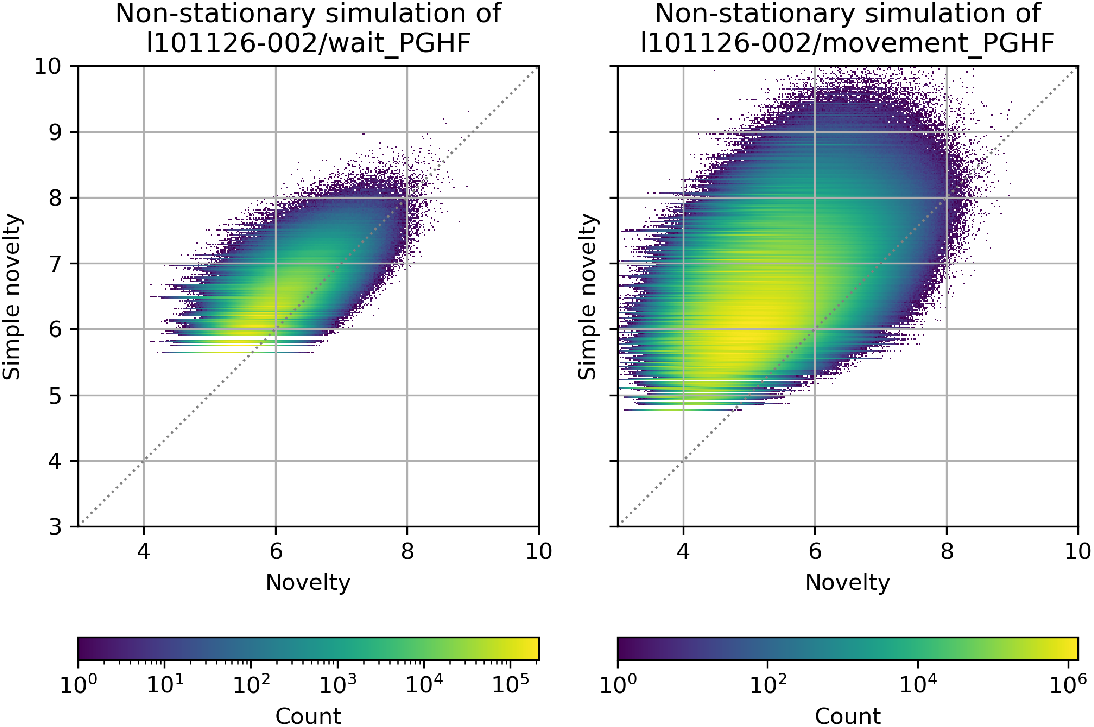
Distribution of simple novelty *sn* against the original novelty *nov*, shown as a pseudo-color 2-dimensional histogram (bin width: 0.02 for both *sn* and *nov* ). Both *sn* and *nov* were calculated for every PR found in 10,000 non-stationary simulations of the ‘movement’ period spike train data to construct the histogram. A brighter color indicates a larger count of PRs with the corresponding combination of *sn* and *nov*, as indicated by the color bar to the right (note the logarithmic scaling of the color code).

### 4.3 Comparison of the quality to its proxies

By extending these observations, the exemplary results in Figs. 7 and 8 encourage the following idea: Our goal is to compare PRs in terms of their ‘quality’ or ‘unexpectedness’. For this, we have several natural quality criteria (e.g., size, repetition, duration, novelty). Here we have defined a canonical ‘ideal’ way of combining these criteria (their quality *Q*). Unfortunately, this measure cannot be calculated directly, but the stationary Quality *Q*_*s*_, which can be calculated, is roughly monotonically related to *Q* and may therefore be used as a ‘proxy’ for *Q* in the comparison of PRs. Once we accept this idea, we could also try to invent other, even simpler proxies for *Q*. This may be very useful in practice, because the calculation of *Q*_*s*_ is still computationally expensive. In the course of inventing more proxies, other than the stationary quality *Q*_*s*_, for quality, we of course also considered the possibility of using the pattern novelty *nov* instead of *sn* in these proxies, because it better captures the real non-stationary dynamics.

Now we formally introduce a few more candidates for such proxies, which either try to approximate *Q* numerically or to show at least a roughly monotonic relationship to *Q*. We can compare and test such proxies by plotting their relation to *Q* for several STPs on the same two multiple-unit datasets, for which *Q* has been estimated by extensive sampling.

Clearly, both the size z and the repetition r of a pattern repetition are relevant, so one could consider combinations of the two. In the course of the so-called SPADE analysis (Quaglio et al. 2017), it became apparent that it makes sense also to consider the duration d of a pattern, but the effect of d is comparatively small compared to that of z and r. Here we have introduced another criterion, namely the pattern novelty *nov*, or the simple novelty *sn*, which directly measures the probability of the PRE and can be used as a replacement for z, but could also be used in addition to z.

Obviously, one can think of many possible ways of combining all or some of these parameters, even without considering additional ones like the precision of pattern epetitions, and any one parameter alone is not sufficient as a quality measure. The effect of d is rather small, and also the effect of combining *nov* and *z* versus using only *nov* is not large, because in the data we analyzed, values of *z* other than 3 or 4 hardly occur. So we have decided to consider only combinations of *r* with one of the parameters *z, nov* or *sn* as different very simple alternatives here, namely *z* · *r*, and three plausible combinations of *nov* (or alternatively *sn*) with *r* .

#### Definition 12

(Quality proxies for a given signature) Given a signature *s =* (*z, r, d, nov* ) we define the following quality measures:

- *S = z* · *r*, which simply counts the total number of spikes that contribute to the pattern repetition event.
- *N = nov* · *r*, which is the pattern repetition novelty (Def. 5).
- 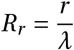, the repetition relative to its expectation,
- 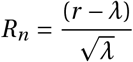, the normalized repetition, where *λ = T* ^*′*^ · *e*^−*nov*^ is the expected number of repetitions.

The first two of these appear quite naturally. However, one can easily create an example where they are clearly misleading. Consider a pattern with a low pattern novelty *nov* (or simple novelty *sn*) which is repeated about as often as expected, i.e., *r = λ*. Such a pattern is not surprising, but still, *S* and *N* can be quite large. This is why we have also introduced *R*_*r*_ and *R*_*n*_ . In the definition of the expected repetition *λ*, one could also use *sn* instead of *nov* . We don’t claim that these quality measures are new; in particular, the last two may have already been used in the literature.

Apart from their extreme simplicity, another property distinguishes these proxies from the computable stationary quality *Q*_*s*_ : this is their ‘locality’, in the sense that the formula requires only parameters from the neurons involved in the pattern, whereas *Q*_*s*_ depends also on the firing rates of other neurons (which it should in principle). This is also what makes the computation quite complex. As long as we are only comparing patterns from the same experiment, we may hope to get a good comparison based only on local properties of the neurons involved. Therefore, we also suggest two more complex local quality measures, which we call the local and the dynamic local quality *Q*_*l*_ and *Q*_*dl*_ . They require the full signature and some more computation, but are still much simpler to compute than *Q*_*s*_ .

In addition to this, we have developed a ‘more dynamic’ version of *Q*_*s*_ which applies a correction *C*_*d*_ to *Q*_*s*_ and is called *Q*_*sd*_ . It allows us to consider the dynamic novelty *nov* instead of *sn* in the formula for *Q*_*s*_ . We realized that this would require some more adjustments, because we essentially assume that all neurons are firing stationarily with a different (typically higher) firing frequency that corresponds to *nov* instead of *sn*. To keep the total number of spikes constant, we redefined *T* ^*′*^ accordingly (see Def. 13 below). This reasoning is described in more detail in Appendix D; it results in the definition of the ‘dynamic stationary quality’ *Q*_*sd*_, admittedly a somewhat contradictory term, which we use in Fig. 8.

These three more complex proxies are defined in the next definition. We have also developed a numerical approximation to *Q*_*s*_ that is based on the Gauss distribution and requires much less computation. It is called the Gauss approximation *Q*_*G*_ and described in Appendix C.

#### Definition 13

(Quality proxies for extended signature) Given an extended signature (*z, r, d, nov, sn*) we define the following quality measures:

- the local quality *Q*_*l*_ *=* − ln (*p*[*R*_*P*_ ≥ *r* ]) − ln(*c*(*z, d* )), where we use the upper bound of Prop. 4, and Prop. 2.
- the dynamic local Quality *Q*_*ld*_ *=* ln(*r* !) − ln(*c*(*z, d* )) − *r* ln(*λ*) − *(T* ^*′*^/*a +* 1 −*r)* ln(1 − *q*) − ln(*rJ+* 1) *+* ln(*r +* 1 − *λ*), where *q =* exp(−*nov* ), *a =* exp((*sn* −*nov* )/*z*), *λ = T q*/*a*. If 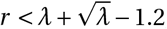, then *Q*_*ld*_ is not evaluated.
- the dynamic stationary Quality *Q*_*sd*_ *= Q*_*s*_ −*C*_*d*_, where *C*_*d*_ is defined (for a fixed value of the size *z*) in terms of the novelty difference *nd* :*=* max(0, *sn* −*nov* ) as follows:

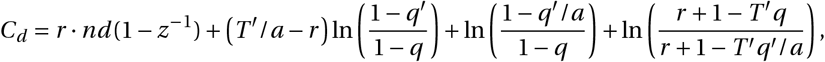

where 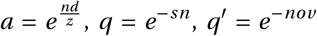. This implies that *C*_*d*_ *=* 0 if *sn < nov*. If *r <* 1.5 · *T* ^*′*^ *q*^*′*^/*a* − 1 then *C*_*d*_ and therefore *Q*_*sd*_ is not evaluated.

In Fig. 10 we display these quality proxies against Q for the *wait* dataset, and in Fig. 11 for the *movement* dataset.

**Fig. 10:**
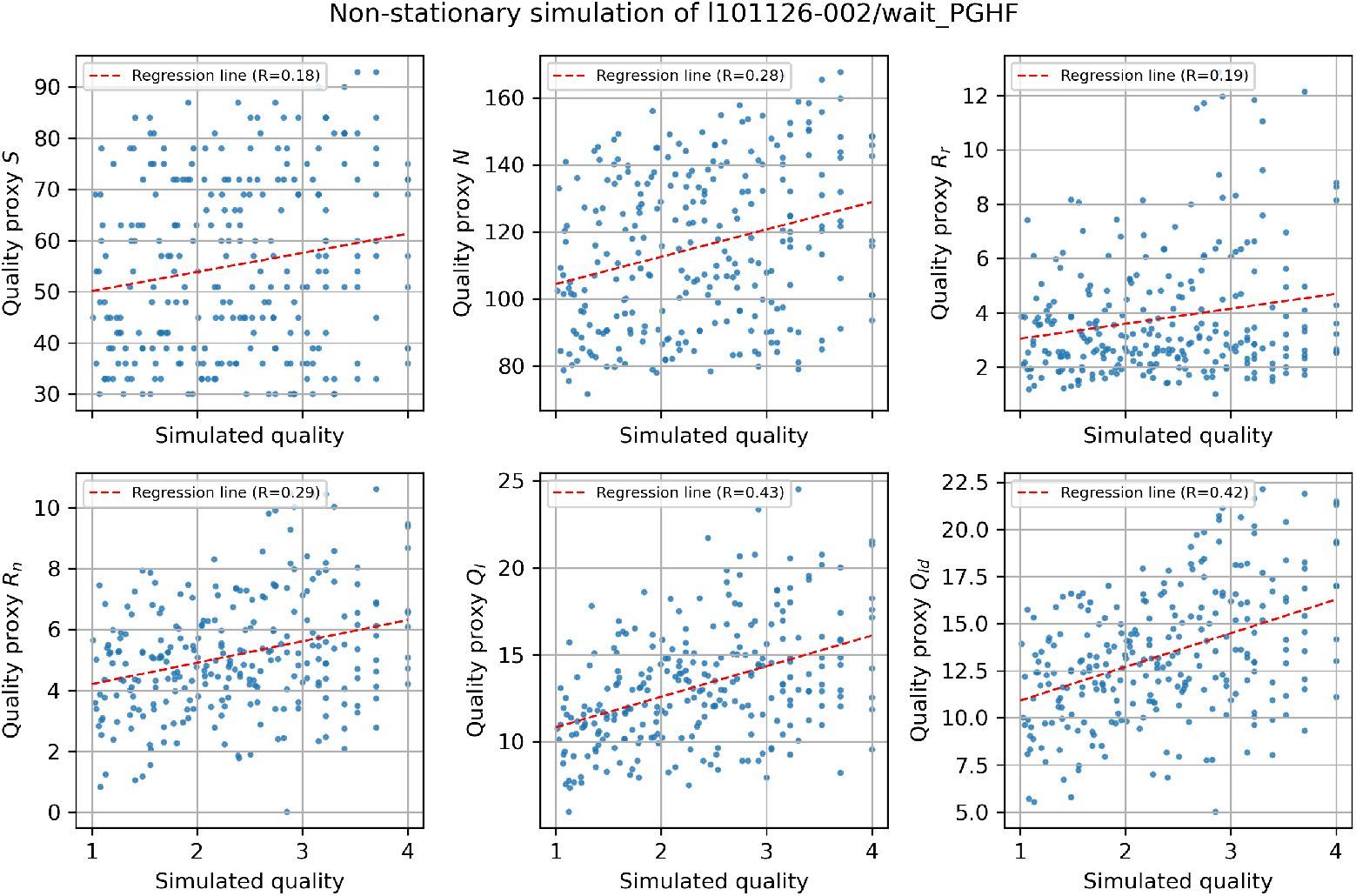
The quality proxies (*S, N, R*_*r*_, *R*_*n*_, *O*_*l*_, and *Q*_*ld*_ ) computed for the ‘wait’ period dataset are plotted against the quality estimate based on non-stationary simulations of the ‘wait’ period (simulated quality, the same data as those used for Fig. 7 (left) and 8 (left)). The red line indicates the least-square regression line, the correlation coefficient *R* of which is shown in the legend.

**Fig. 11:**
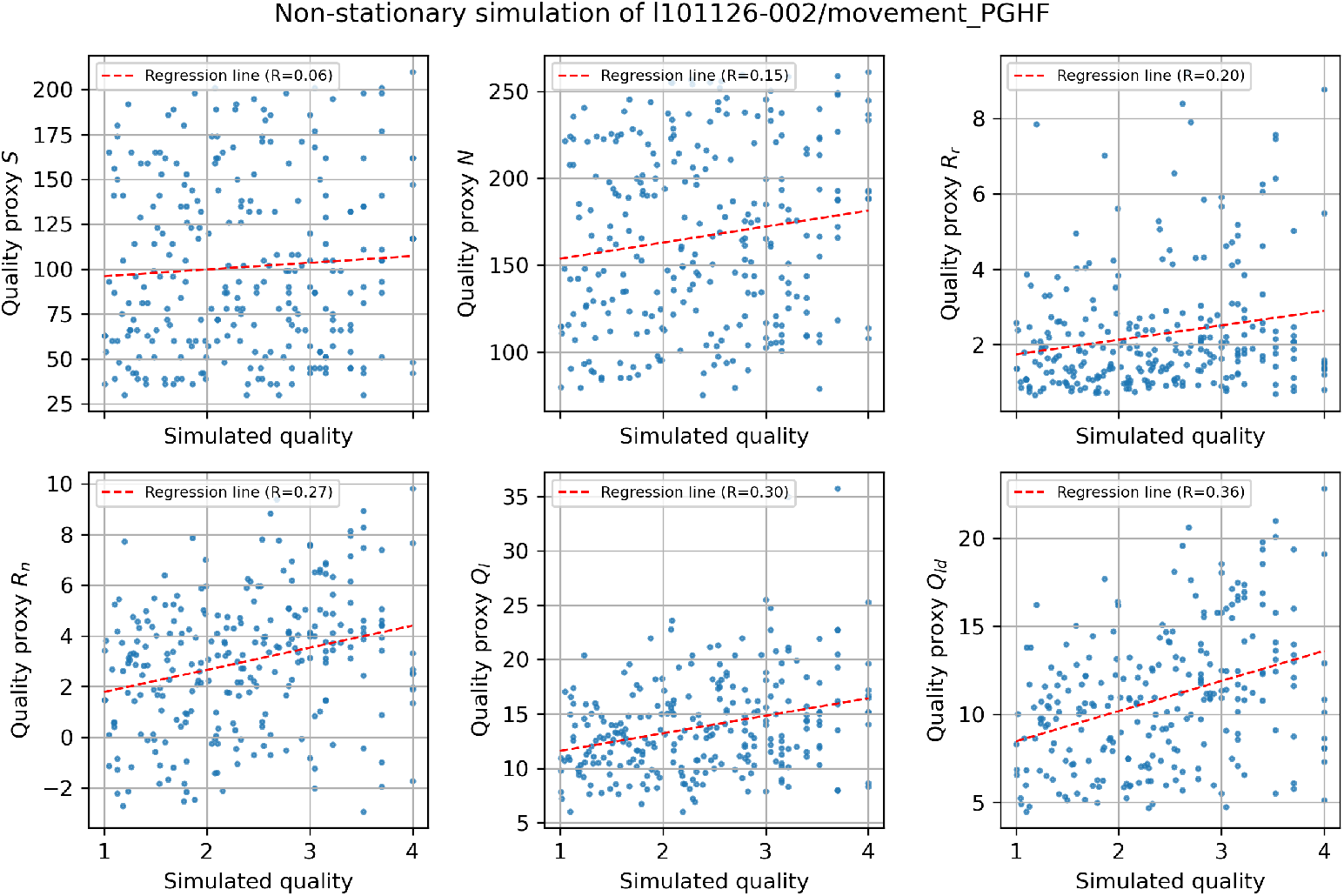
Same plots as in Fig. 10, but computed for the ‘movement’ period dataset.

**Fig. 12:**
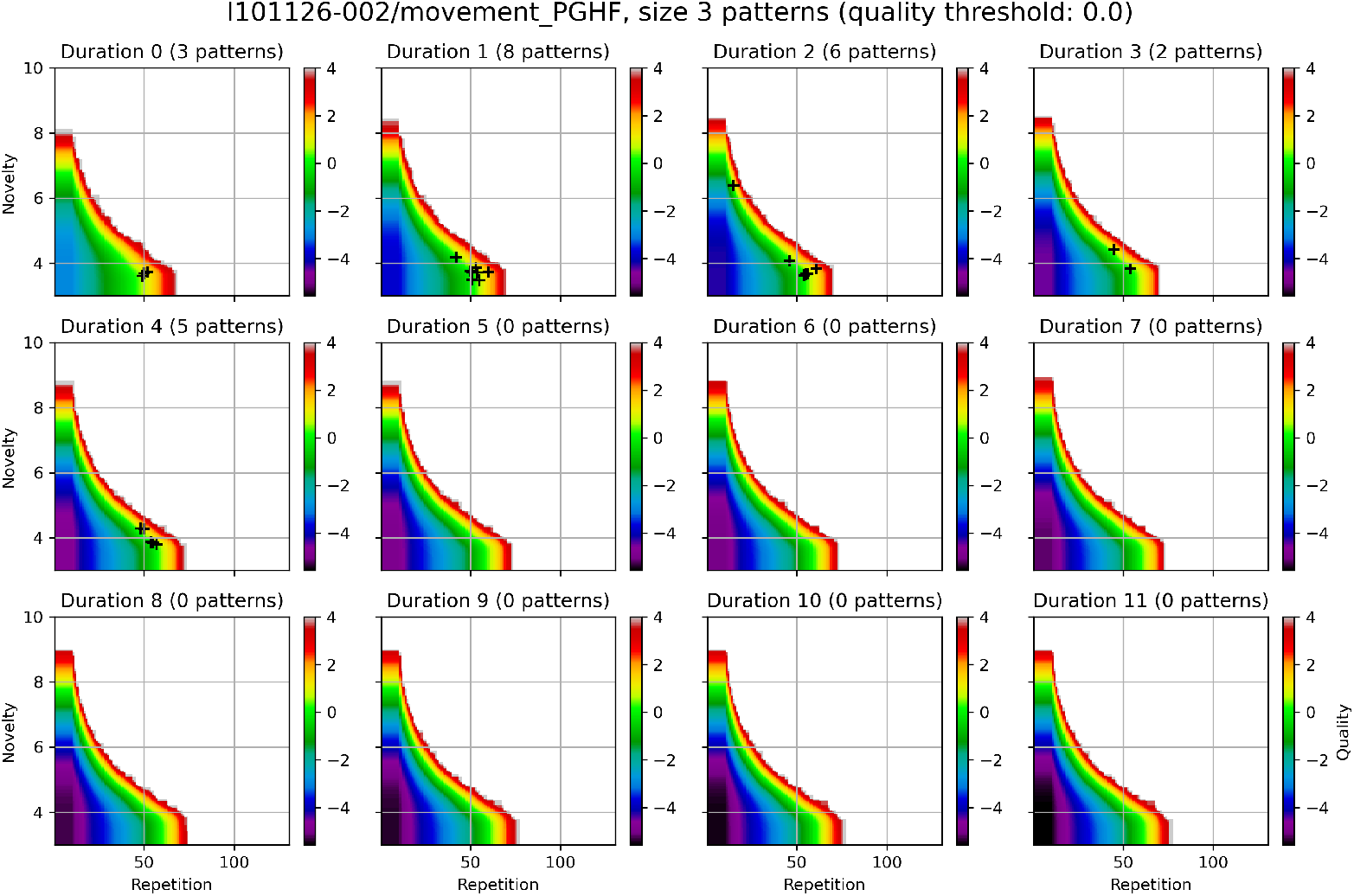
Signatures of STPs found in experimental spike trains displayed in the null hypothesis quality spectrum. Each panel shows a 2-dimensional slice (spanned by repetition and novelty; size is fixed to 3) of the 4-dimensional quality histogram estimated from 10,000 non-stationary simulations of the ‘movement’ period dataset, with black crosses indicating the signatures of the PRs found in this dataset with quality values greater than 0.0. Quality values are represented by colors as indicated by the color bar to the right; the region of no color represents the signatures that are never observed in any of the simulations (and hence the quality cannot be estimated).

Taken together, we have defined here 9 proxies for our defined quality, namely *S, N, R*_*r*_, *R*_*n*_,*Q*_*l*_,*Q*_*ld*_,*Q*_*sd*_,*Q*_*s*_,*Q*_*G*_ . All these proxies share with the real quality the basic monotonicity property (with respect to the signature parameters) and they all correlate more or less with the real quality *Q* that we defined in Section 2 (and that was estimated for the two datasets from 10,000 simulations).

Fig. 11 shows that for a given collection of quality criteria in a signature, different combinations of these criteria can result in quite different quality proxies that only roughly point in the same direction. Based on the regression coefficients the lower three of these could probably be used in data analysis, i.e. *R*_*n*_,*Q*_*l*_,*Q*_*ld*_, if one wants to avoid more complex calculations. Different collections of criteria can also be considered. In particular, one could consider a precision criterion, which bounds the amount of jitter in the delays between subsequent spikes in a pattern when it reoccurs at different times. Presently, the maximal jitter is determined implicitly by the bin size Δ*t* . Our new criterion of pattern novelty certainly reflects, to a large degree, such a precision criterion; similarly, it also reflects the size of a pattern. It may therefore be considered a matter of taste whether one wants to include the size and/or some precision criterion in addition to the pattern novelty in the signature, if one is particularly interested in, for example, finding large patterns or very precise repetitions. Practically, one can use some reasonable quality proxy for an easy direct comparison of PRs, and then focus on the top 10 or 20 PRs for closer analysis of further quality criteria or, more importantly, their occurrence times in relation to the behavioral context.

## 5 Using Quality for Hypothesis Testing

For a statistical test one needs at least a ‘test variable’ or ‘test statistics’ *T* and a ‘null hypothesis’ that determines the corresponding ‘null (hypothesis) distribution’ of *T* . In the early days of statistics, the test variable *T* was usually obvious, and its null distribution could be easily calculated from simple statistical assumptions. Over the years several more statistical tests were designed and the corresponding null distributions were calculated and presented in statistics books. With the increased speed of computers, this situation has changed drastically. Nowadays, everyone can basically create their own statistics by defining some test variable, *T*, which appears to be appropriate, without needing a mathematician to work out the corresponding null distribution. This distribution, even if it is not exactly defined mathematically, is then determined implicitly by extensive sampling, i.e., by a procedure that generates sample values of *T*, also called surrogate data.

In the statistical analysis of multiple-unit spike patterns, the problem that we have discussed here is the definition of the most appropriate test statistic *T* that combines several plausible quality criteria into one variable. For this purpose, we want to use the pattern quality *Q* or one of several simpler proxies for it. Intuitively, the problem is to find the right direction in the 4- or 5-dimensional space of quality criteria that best captures our idea of ‘quality’ of a PR. It is important to realize that, whatever we choose for *T*, combined with an appropriate null hypothesis, it will create a valid statistical test. It may only be suboptimal for our purposes if it does not point in the right direction.

Unfortunately, we cannot directly use *Q* as a test statistics, since it is not defined as a random variable defined on the MUST process. The easiest and perhaps most intuitive way to turn it into a random variable is to define the quality *Q*(*x*) of a MUST realization *x* as the maximal quality of all PRs found in *x*. This, however, is a very strict definition. It is also possible and plausible to consider ‘weaker’ definitions of the test statistics, in particular, the top-k quality *Q*_*k*_ (Def. 7).

The other ingredient, the choice of a proper null hypothesis, is also a delicate issue. In the context of neural coding, in particular when it concerns the temporal precision of single spikes versus their determination in terms of the much lower temporal resolution of firing rates of the participating neurons, one usually relies on the generation of surrogate data for the null-hypothesis. The idea is to destroy any potential higher temporal precision by ‘jittering’ or ‘dithering’ individual spikes (or groups of consecutive spikes) by a small temporal amount which would only minimally change the firing rates measured at a much lower temporal resolution. This method of ‘dithering’ corresponds to a ‘null hypothesis’ which is very close to the original data and therefore not easy to falsify.

In principle, one could define a stochastic model of this jittering process, which could then be considered as a statistical null hypothesis, but this null distribution would require as parameters all the actual spike times of all recorded neurons. Being trained in classical statistics, where typically only a few parameters are needed to specify the null distribution, most of us would feel uneasy with such a definition, and it would also be practically impossible to use it for mathematical calculations. But it can still be used as an argument to justify such a method as a proper statistical test, if such an argument appears to be necessary. Indeed, versions of jittering have been proposed that require much fewer (but still many) parameters from the experimental data and that can be formulated more naturally as a statistical null hypothesis (Amarasingham et al. 2012). In a way, our initial definition of the MUST process is also of this type.

Here we propose to use the stationary quality *Q*_*s*_ (or rather the much faster computable Gauss approximation), or perhaps the dynamic stationary quality *Q*_*sd*_, in Def. 7 to generate a test statistics to determine the ‘significance’ of a PR, as being unlikely to be generated under the null hypothesis, and the simulated MUST or some version of dithering or jittering as the null hypothesis. We have to realize that with this statistics we are effectively comparing the experimental STPs with the best (or the k best) randomly generated ones.

In practice, the following steps are needed according to our proposal to find potentially interesting repeating spike patterns in multiple-unit recordings and to evaluate their statistical significance:

1. Decide on a bin-size Δ*t* and bin the spike trains into binary bins. In our example datasets, we used Δ*t =* 5 *ms*.
2. Find interesting pattern repetitions, for example, using frequent itemset mining (Borgelt 2012).
3. Choose a proxy *Q*_*x*_ for the quality of patterns, for example, the stationary quality or the Gauss approximation.
4. Calculate the values of *Q*_*x*_ of the pattern repetitions found. For this, we need their signature.
5. Choose a null-hypothesis, i.e., a dithering method or the MUST simulation.
6. Create many surrogate or simulated data.
7. For each surrogate dataset, use Def. 7 to compute the corresponding value of the test statistics *T* . Arrange these values to create the ‘null distribution’ of *T* .
8. From this distribution read off the p-values of the quality of the patterns found in the original data.

When we list all the necessary steps like this, the procedure appears quite complicated, involving several decisions to be made until one arrives at a statistical test. However, given the flexibility of defining and implementing statistical tests today, this situation is quite common in modern data-intensive experimental science. There are many ways of reorganizing and transforming data, and many practical decisions to be made, before one can perform a statistical test. Usually, just this test and its result are presented, and decisions are made on the way, and potential other test options are just not mentioned.

This is also the reason why we don’t present test results yet in this methodological paper.

Instead, in Fig. 12 we illustrate one potential alternative way of analyzing experimental data by displaying the signatures of spike patterns found in the data in the quality spectrum estimated based on simulations or surrogates. Here the signatures of the PRs found in the ‘movement’ period dataset with high quality values (here *Q* ≥ 0.0) are indicated in (2-dimensional slices of) the 4-dimensional quality spectrum estimated based on non-stationary simulations as explained in Section 3.2. One can see that a majority of the high-quality PRs are with rather low novelty values, indicating that those patterns are composed of neurons with high firing rates. However, there is one pattern with a relatively high novelty value (in the slice of duration 2), meaning that the neurons composing this pattern have low firing rates and hence they should be a separate group of neurons from those composing the other patterns with the low novelty values. Thus, with this type of display, one can gain a concise overview of the statistical properties of potentially interesting STPs found in the given dataset, without performing significance tests.

## 6 Discussion

Our main contribution in the present study is a new criterion for the evaluation and comparison of spike pattern repetitions or repeating spike patterns in multiple neuron spike-trains. We are not just adding one more criterion to the existing ones, but we provide a canonical and conceptually simple way of combining any number of quality criteria into one. Here we want to answer the following questions: Why and for whom are we doing this? What are our additional contributions to this community? What are our basic assumptions, and to what extent can they be loosened? Are there closely related problems that we did not consider? Why are we interested in spike patterns in the first place?

### 6.1 Why spike patterns?

There are two converging motivations, which come from the interest in neural representations, sometimes also called neural coding (Perkel and Bullock 1967), and from the interest in neural computation, here in particular the question of how the observed neural responses are possibly produced by the interactions of the many neurons that lead up to the neuron that has been recorded.

The first interest motivates most of classical neuroscience, where the activity of the recorded neuron(s) is related (or correlated) to the events in the outside world, i.e., the stimuli and responses of the experimental paradigm. Here, there was a long-standing debate about whether to consider representations by neuronal firing rates or by the precise timing of individual spikes. Obviously, the exact spike times contain more information than the averaged rates, but do they also contain more information about the experimental stimuli and responses?

In the usual experimental paradigm, the animal was confronted many times with the same stimulus or task, and the neuronal responses were accumulated across these repetitions in a so-called PSTH (Gerstein and Kiang 1960). This type of averaging gave the impression that the individual spikes appeared much less regularly than their accumulated firing frequencies, which co-varied with the stimulus or the animal’s response. However, if the animal has to react quickly and correctly to the stimulus, it cannot wait for repetitions or for temporal averaging. So the idea came up that perhaps the next level of computation is based on spatial averaging across neighboring neurons that react roughly in the same way to the stimulus. Computationally, this seems to be quite an inefficient use of neurons, and therefore, it seems to be reasonable to look for additional sources of information beyond averaged firing rates, in particular, in the more exact timing of spikes. Since there is no external clock, this of course means the timing of spikes across several neurons relative to each other.

Also, beginning with Donald Hebb’s classical book (Hebb 1949), functional brain models have always relied on representations in terms of temporally co-active groups of neurons, in particular in cell assemblies or synfire chains (Palm 1982; Abeles 2012). However, these theories are primarily based on particular associative connectivity structures, which are presumably formed by synaptic plasticity (auto- and hetero-associations, see Palm (2013)) and with respect to parameters like the size or the expected number of activated groups they are not sufficiently concrete to make definite predictions concerning spike patterns (Palm 1993; Harris 2005; Kleinjohann et al. 2024).

One way of showing that the precise timing of spikes is not just random, but can be used in the brain, could be the observation that precisely timed spike patterns across several neurons actually do occur repeatedly. More exactly, they should occur more often than expected by ‘chance’, where chance means the null-hypothesis, which should include the stimulus-evoked variations and covariation in neural firing rates that we know from classical neuroscience. This results in statistical tests based on a non-stationary null-hypothesis, as our MUST, or on surrogate data generated by spike dithering or jittering methods (see Stella et al. (2022)). This high sophistication in generating the null-hypothesis is necessary if we want to show that spike-patterns can contain more information about the situation beyond what can be inferred from the firing rates, and of course, it makes the job of finding interesting pattern repetitions much harder.

In order to understand the computation in networks of neurons, one has to study their interaction. This can be done by pairwise correlations between neuronal spike-trains, but then it may also become necessary to study higher order correlations, because one single input spike is not enough to fire a neuron, and it usually needs the convergence of several input spikes, which is even more effective when they arrive at the same time (Abeles 1982). Higher-order correlations have been measured, but this quickly becomes very cumbersome if one goes beyond third order (Palm and Pöpel 1985; Staude et al. 2010a). The fact that spikes occur sparsely can help here, because the occurrence of a spike provides much more information than its non-occurrence.Therefore, one may look for co-occurring spikes, i.e., spike patterns across multiple neurons.

The search for coincident spike patterns has been quite successful (Riehle et al. 1997; Grün et al. 1994, 2002; Maldonado et al. 2008; Kilavik et al. 2009; Pipa et al. 2013; Torre et al. 2016a; Shahidi et al. 2019) but in view of variable traveling times for spikes from the cell body to the synapse and from the synapse to the cell body of the postsynaptic neuron, coincident arrival on one postsynaptic neuron may not mean coincident recordings of those spikes at the cell bodies. This has led to the idea of polychronization (Izhikevich 2006). In addition, longer-lasting coincident activity may stay active by moving from neuron group to neuron group in so-called synfire chains (Abeles 1991, 2012). For these reasons, it may be necessary to look not only for coincident patterns, but also for temporally (and spatially) extended multi-neuron spike patterns. Unfortunately, this increases the number of potential patterns considerably, which complicates the statistical analysis.

### 6.2 Why quality?

Searching for ‘interesting’ spatio-temporal spike-patterns is problematic because there are so many potential patterns. Indeed, if every single spike counts and there are no particular a priori requirements on patterns to ‘look nice’ or ‘beautiful’, then essentially every set of spikes can be called a pattern, and the number of potential patterns can grow exponentially with the number of spikes. The only, or at least the first criterion to select patterns and reduce this huge number, is their repetition. Another obvious criterion is their size, sometimes also called ‘complexity’. For example, one can require a pattern to be repeated at least twice to be considered potentially interesting. In this paper, we have argued that the size should be at least 3 in order to call it a pattern. So it has become commonplace to put lower thresholds on the size and number of occurrences of the patterns to be searched for. Using these two parameters, one can practically control the total number of patterns to be considered further. In the related area of ‘frequent itemset mining’, the same two parameters are also used for this purpose. In neuroscience, the dimension of time also plays an important role, which again increases the number of potential patterns and adds two more parameters that can be used to control it, namely the duration of a pattern and the precision of its repetitions. Finally, we can also consider the probability of a spike to occur, the logarithm of which we used to define the pattern novelty. In most studies, several of these parameters are considered, almost always at least two. If these parameters are pushed very high, one can end up with very few candidate patterns for further analysis. But which parameters should be pushed how high?

The statistical solution to this problem would be to construct a significance test for patterns and consider only patterns that are significant below a given significance probability threshold. This does not work, however, when one starts with a large number of patterns, or rather pattern repetition events, because one is faced with an enormous multiple testing problem, which cannot be tackled by the usual methods. Fortunately, this problem can be reduced quite drastically if one considers all patterns with the same signature as equivalent and only constructs a test for these signatures. This was one of the initial ideas in the SPADE project (Torre et al. 2013). The signature contains the parameters mentioned above, and there are still at least two, in the example considered here, there are three. This implies that there is still a considerable (although much smaller) multiple testing problem. In the early days of the SPADE project (Torre et al. 2013), the testing of signatures instead of individual patterns reduced the multiplicity of tests from millions (the exact numbers can be calculated from Prop. 2) to about one or two hundred (the number of entries in the so-called p-value spectrum). In fact, when we take the repetition of a pattern as a test variable, the number of tests can be reduced to just a few for the different sizes, but after adding the duration to the signature, it increases to about a hundred. This situation becomes much worse when we also add the novelty to the signature, which is a continuous variable, but it is very useful to consider the novelty since it avoids overestimating the quality of patterns in rapidly firing neurons.

To avoid the multiple testing problem altogether, it would be desirable to have just one parameter to control the number of candidate patterns. This is why we introduce our quality measure, which combines the quality criteria collected in the signature into one, and also directly controls the number of remaining candidate patterns. The idea is simply to use the different quality criteria to define a partial order relation ≤ for the comparison of patterns. We then use the cone based on a single signature (see Fig. 3) to estimate the number of pattern repetitions in this cone. This number, the ubiquity of a signature, is used for our quality measure. Based on this, we can not only compare repeating patterns directly, but also define a test statistic for significance testing that can also be applied to individual patterns or their signatures.

### 6.3 Our basic assumptions

Our mathematics is based on the definition of the MUST. Here, we have decided to use discrete instead of continuous time. And we have assumed independence of the variables 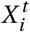. This is essentially the simplest stochastic model of a non-stationary multi-unit spike-train. We could have formulated the model in continuous time, but then our independence assumption concerning the time index *t* would be too strong, and we would probably use some more complex parametric model like the gamma-process for the spike trains. Also, exact repetition of a pattern would have probability zero; therefore, we would introduce a precision criterion, for example, requiring that the temporal jitter in delays *d*_*i*_ between successive pattern spikes across all occurrences of the pattern has to be less than a threshold *δ*. Then our definitions would also make sense, but for the calculations, we would have to require a minimal separation (in time) between successive occurrences (Def. 4) to argue that the corresponding successive spikes in each of the pattern neurons are almost independent. Then we could essentially use *δ* instead of Δ*t* to compute the basic time-dependent firing probabilities 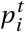. The whole argument would become much more clumsy, and, anyway, with our MUST model, we are getting close to it for a small bin-size Δ*t* . Actually, our definitions of ubiquity and quality make sense and are potentially useful not only for neuroscience, but for any case of temporal repetitions of binary ‘patterns’ or sets, as in frequent itemset mining, which can be described by finitely many binary random variables 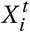. The statistical problems concerning the large number of patterns and multiple testing are essentially the same (Kirsch et al. 2012). The ‘pattern’ aspect is described in the most general way by a preorder ≤ that may involve several criteria, potentially also ‘beauty criteria’.

### 6.4 Aspects we did not consider

In our mathematical analysis, some things are simply taken for granted, in particular the bin-size Δ*t* and the probabilities 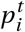 of having a spike in a bin. For the experimentalists, both of these are delicate issues. In order to determine the parameters 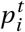, one has to estimate the so-called instantaneous firing rates from the experimental spike trains. Unfortunately, there is no obvious simple solution to this problem, because the method depends on the purpose for which you need these values; this is discussed in quite extensive literature, see for example (Koštál and Kováčová 2025; Tomar 2019).

By taking too large bins Δ*t*, we obviously lose precision in the timing, and we also lose spikes, because it will happen more often that more than one spike is in the same bin. By taking too small bins, we are expecting a high precision in the repetition of a pattern, and also we will more often miss pattern occurrences because one spike is not found in the prescribed bin, but in the next one. In addition, the assumption of independence between neighboring bins of the same neuron is more unrealistic. This problem can be alleviated by introducing a larger minimal separation between subsequent occurrences of the same pattern. Also, the first problem can be solved by allowing inexact pattern repetitions and introducing a precision criterion, i.e,. a threshold *δ* for the maximal allowed jitter in pattern inter spike intervals (see above) that is larger than Δ*t* . This can be easily defined, but increases the complexity of finding those pattern repetitions considerably (Borgelt 2012).

Actually, we also didn’t talk about methods of searching for pattern repetitions. We consider this a separate issue, and we cited a few of these methods. Often, the evaluation of patterns is intermingled with the search process because one does not want to collect and store too many uninteresting patterns, but conceptually, we prefer to separate these two problems. In SPADE, we used FIM (frequent itemset mining) for the search, which appears to be one of the fastest methods, but it uses binary patterns and discrete time. It also requires exact repetitions (i.e. *δ =* Δ*t* ).

Conversely, when using a low Δ*t*, we could post-process the patterns found by calculating their precision *δ* which will be smaller than Δ*t* and could also be used to recalculate the quality of the pattern.

Another thing we take for granted is the number *N* of ‘recorded’ neurons. It will often occur that one finds one or several repeating patterns in just a few of the recorded neurons. In such a case, it may be tempting to consider only these neurons as the recorded ones, because it simplifies the calculations and gives higher quality values. One can perhaps use these quality values for comparison within this experiment, but statistically, this would not be correct. Something similar might sometimes be done: often, the patterns occur in those neurons that react with high firing frequency to the stimulus. So one could set a threshold on the (average or maximal) firing frequency and discard all neurons below that threshold from the ‘recording’ because they are ‘not reacting’ or ‘not involved in the computation’.

### 6.5 What else?

Conceptually, our quality measure may be quite simple and plausible, but unfortunately, it is not easy to compute or estimate, and the calculation requires strong assumptions (stationarity of spike-trains under the null hypothesis). In fact, the estimation of quality values from simulations or surrogates requires a huge number of them, because we are interested in very rare events; for the quality spectrum in Fig. 12 we used 10,000 simulations and FIM runs, where FIM found billions of patterns, and even this was hardly enough, 100,000 would have been better. For the calculation of the simplified formula in Thm. 6, we had to calculate and sum more than 540,000 small probabilities. Therefore, we consider the quality measure as a mathematically defined ideal combination of quality criteria, which may practically be replaced by a ‘proxy’ that is also more or less plausible and much easier to calculate. Since not everybody has a supercomputer and there will be much data to evaluate (in particular, many pattern signatures found in a few hundred surrogates), we propose a few simpler proxies for quality that seem to correlate quite well with it. Up to now, we only tested this on two experimental datasets, so this line of research could be extended in the future.

The use of a proxy for quality that can be directly calculated has another great practical advantage. You do not need to simulate the MUST process. And if you use a proxy that is based on the simple novelty *sn* instead of the novelty *nov*, you don’t even need the ‘bin firing probabilities’ 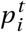. You can also use the proxy for a statistical test combined with your favorite 0-hypothesis or surrogate method as described in Section 5. We are now preferring a surrogate method called ‘trial shifting’ (see Stella et al. (2022)), but we have not yet decided which test statistics to use (we are tending towards the top-k statistics (Def. 7) with a small value of *k*, e.g., 1 - 4, for this experiment).

Essentially, we have used the MUST as a framework to define all our quality criteria, and finally, the ubiquity and quality of a repeating pattern or its signature. Then we used the stationary MUST to get a feeling for the size of these quantities and finally to compute them. Based on this, we suggest using related similar proxies for comparison and testing of PRs, which is much simpler and can be combined more easily with methods for pattern search.

1

## Statements and Declarations

### Availability of Data and Materials

The data and codes that support the findings of this study are available in Zenodo with the identifier: https://doi.org/10.5281/zenodo.18424512.

### Author Contributions

Conceptualization: G.P., S.G., A.S, J.I.; Data curation and validation: J.I.; Formal analysis and investigation: M.P., G.P., J.I.; Funding acquisition and resources: S.G.; Methodology: G.P., M.P., J.I.; Project administration: G.P.; Software: J.I., G.P., M.P.; Supervision: G.P., A.S., S.G.; Visualization: J.I., M.P.; Writing - original draft preparation: G.P., M.P., J.I., S.G.; Writing - review and editing: G.P., M.P., J.I., S.G., A.S.;

## Acknowledgments

The authors express their gratitude to Peter Bouss, Jonas Oberste-Frielinghaus, and the Statistical Neuroscience Group of IAS-6 for stimulating discussions. This study is partly funded by NRW-network ‘iBehave’ (grant number: NW21-049) and the Deutsche Forschungsgemeinschaft (DFG, German Research Foundation) - grant 368482240/GRK2416, grant 561027837/ GR 1753/9-1, and grant 491111487 (Open Access Publication).

## Appendix A Proof of Proposition 5

Here we show the following estimate of the tail *B* (*q, T* ^*′*^, *r* ) of the binomial distribution:

If 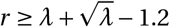, it holds that

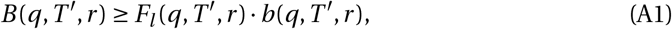

where

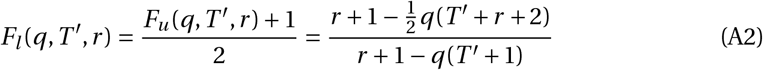

*Proof* Fig. A1 shows that *a*_*i*_ ≤ *b*(*r + i* ), *a*_0_ *= b*(*r* ), *a*_1_ *= b*(*r +* 1), and *a*_*i*_ *= b*(*r* ) − *i* · *d* with *d = b*(*r* ) − *b*(*r +* 1). Also, it holds that *d* : 1 *= b*(*r* ) : (*k +δ*). This shows that

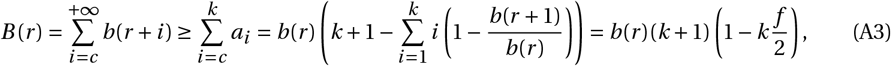

where 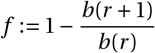.

**Fig. A1:**
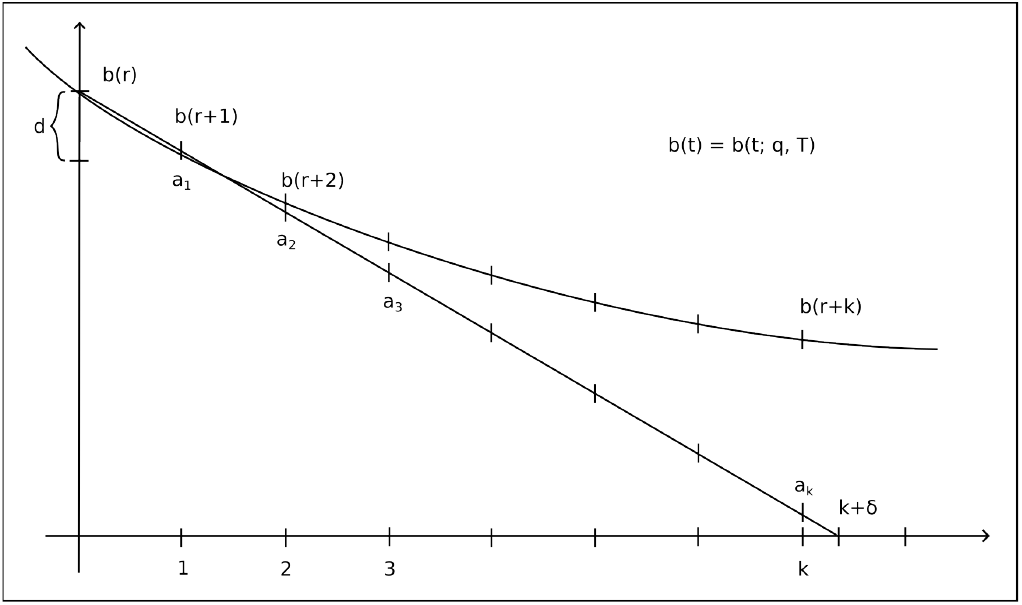
Schematic representation of the inequalities employed in the proof.

Now 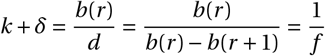. Combining this with (A3) yields

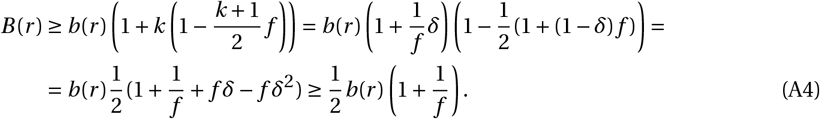

We observe that 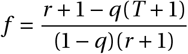 and therefore 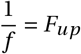 from Prop. 4.

Finally, the drawing in Fig. A1 requires that *b*(*r* ) ≥ *b*(*r +* 1), i.e. that *r* ≥ *λ* :*= T q*, and that *b*(*r* ) − *b*(*r +* 1) ≥ *b*(*r +* 1) −*b*(*r +* 2), i.e.

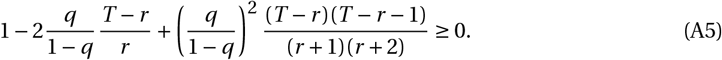

Defining *x = r* −*λ* we obtain from this a quadratic equation for *x* in terms of *λ* and *q*, namely

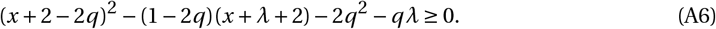

Solving this equation, we obtain 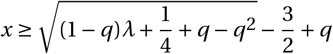, i.e.

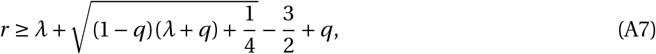

which is the restriction on *r* for the lower bound to hold. Since this condition is somewhat clumsy, we can use a slightly stricter, but simpler condition, which works for *λ* ≥ 1, namely 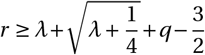. Remember that we are considering repetitions, i.e., we assume *r* ≥ 2 anyway (which also works for *λ* ≤ 1). In the practical calculations, we may use an even simpler condition: 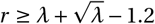, which works for *q* ≤ 0.1.

## Appendix B In Theorem 6, *G*_*a*_ (*z*) decreases fast with *z*

We first observe that for any reasonable value of *nov* or *sn*, we get 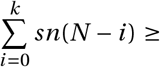 *nov* for some typically small value of k (here we assume, as in the theorem, that the neuron indices are ordered with increasing firing rates). This implies that for *z* ≥ *k* the condition that −*log* (*q*) ≥ *nov* is automatically fulfilled and we don’t need the indicator functions 𝟙_*j*_ in the sums of *G*_*a*_ (*z*). In this case, we look at the quotient 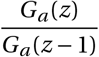 and see that in 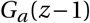 the last sum of *G*_*a*_ (*z*) has essentially been left out and in the last term (1−*q*) ^*T′ −r*^ *F*_*a*_ (*q, T* ^*′*^, *r* ), *q* is replaced by 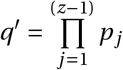 If we compare this with the next quotient 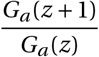, we can observe that 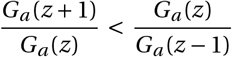 and we can stop the summation, for example, when 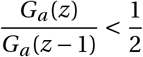. Practically, we rely on this argument in our program to compute the utility.

Since this argument may not be completely obvious, we also give here another, very rough estimate of this quotient, which still shows that it will be small for reasonably large k. In the summations of *G*_*a*_ (*z*) we now assume that the neuron indices are ordered by decreasing firing probability, and we estimate the last sum (for different values of the previous indices) by its largest possible value. This is

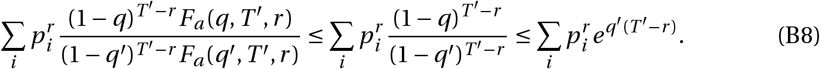

With this we obtain 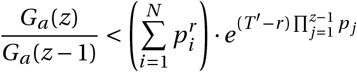.

To see that this becomes very small for not very large z, we observe that for 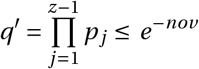 and assuming that 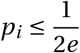 (for *i* ≥ *z*) we obtain

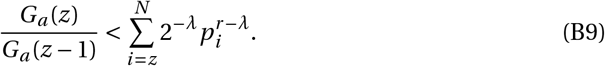

The example below illustrates that this estimate is really small for realistic firing probabilities and signature values.

## Appendix C Gaussian approximation of the ubiquity

We noted that computing the term *G*_*a*_ (*z*) requires significant computational effort, as the number of terms in these sums grows almost proportionally to *N*^*z*^ . Therefore, performing the full sum over all values of z up to N is not practical. Fortunately, we observed that after a few summations (typically up to z=5 or 6), the values of *G*_*a*_ (*z*) decrease exponentially, allowing us to truncate the summation without significant loss of accuracy. The reasons for this rapid decay are further elaborated below.

Since the evaluation of the term *G*_*a*_ (*z*) in the formula for the ubiquity in Thm. 6 is computationally very complex, we have also tried to avoid this computation and to approximate it by a Gaussian integral. The resulting approximation of *Q*_*s*_ is called the Gaussian approximation *Q*_*G*_ . The idea is roughly the following: It is quite common to assume that average firing rates in neural populations are log-normally distributed (Roxin et al. 2011). If this is the case for the population of neurons from which the MUST has been recorded and if the firing rates are low, then one may assume that the simple novelties 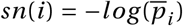 of the single neurons are normally distributed and also the pattern novelties *sn*(*P* ). Then one can compute *G*_*a*_ (*z*) by a Gaussian integral.

So we can fit the simple novelties *sn*(*i* ) of the recorded neurons in the experiment by a Gaussian distribution and obtain the mean *µ* and variance *σ*^2^ of this Gaussian distribution *g*_*z*_ of *sn*(*P* ) across all patterns *P* of size z.

Based on this, we get

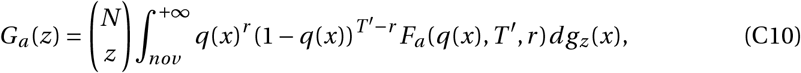

where *q*(*x*) *= e*^−*x*^ . Let us call *H*_*a*_ (*q*) *=* (1 − *q*(*x*))^*T*^ *′*^−*r*^ *F*_*a*_ (*q*(*x*), *T* ^*′*^, *r* ). Then, we have

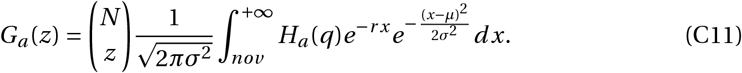

Moreover, naming *µ*^*′*^ *= µ*−*rσ*^2^, we have

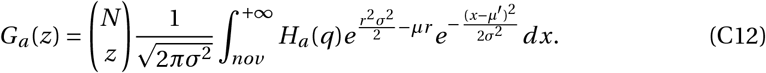

Finally, let us call 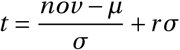. Changing the variable of integration, we have

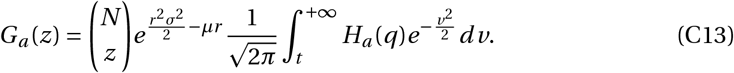

This could be easily evaluated by the standard Gaussian integral, if *H*_*a*_ (*q*) did not depend on *v* (via *q*). Fortunately, it only varies slowly with *v* and increases towards 1 as *v* increases to infinity. Therefore we can split the interval [*t*, ∞) into a few smaller intervals in which the term *H*_*a*_ (*q*) does not change too much and calculate new upper and lower bounds for the integral on each interval using *F*_*u*_ and *F*_*l*_.

## Appendix D Reasoning behind the proxies *Q*_*sd*_ and *Q*_*ld*_ in Def. 13

The idea for these proxies is to use the real novelty *nov* instead of the simple novelty *sn* in the formula for *Q*_*s*_ .

Usually, spike patterns are occurring at time periods when the firing frequencies of almost all neurons are evaluated. For this reason *sn* is typically larger than *nov* . Assuming stationarity would imply evaluated firing frequencies during the total time *T*, which would be inconsistent with the given firing probabilities *p*_*i*_ . To resolve this at least roughly, we assume in these definitions that all neurons are firing with elevated frequencies during a shorter time. This means that we replace *p*_*i*_ by *p*_*i*_ · *a* and *T* ^*′*^ by *T* ^*′*^/*a*, where *a =* exp((*sn* − *nov* )/*z*).

In this situation we don’t need to redo the calculation of *U*_*s*_ for the modified values, but we can just compute a correction term *C*, based on the quotient of the modified and the original value of *U*_*s*_ . The correction is only applied if *sn > nov* . Also, a too small value of *nov* may cause a very large upper bound for the binomial tail and violate our condition for the lower bound. Therefore we require a rather strong condition on *r* in order to evaluate *Q*_*sd*_, namely *r <* 1.5 · *T* ^*′*^ *q*^*′*^ − 1 (see Def. 13). This correction has to be applied for each fixed value of the size *z*.

Essentially the same correction is done for the definition of *Q*_*ld*_ from *Q*_*l*_ . Here we use a slightly weaker condition in order to evaluate *Q*_*ld*_, namely the condition for the validity of the lower bound in Prop. 5.

## Appendix E Signatures used in Figs. 6, 7 and 8

Here we show the signatures used for the plots in Figs. 6, 7 and 8 in terms of combinations of size *z*, repetition *r*, duration *d*, and simple novelty *sn*.

For the comparison of the computed stationary quality to the quality estimate based on stationary simulasions (Fig. 6), we used the following signatures:

- For the “wait” period dataset (*z* is fixed to 3; *sn* is varied in steps of 0.1):
  - 11 ≤ *r* ≤ 19, *d =* 6, *sn =* 7
  - 12 ≤ *r* ≤ 22, *d =* 6, *sn =* 6.4
  - 18 ≤ *r* ≤ 28, *d =* 6, *sn =* 5.8
  - 20 ≤ *r* ≤ 32, *d =* 6, *sn =* 5.8
  - 20 ≤ *r* ≤ 32, *d =* 6, *sn =* 5.8
  - *r =* 15, 0 ≤ *d* ≤ 11, *sn =* 7
  - *r =* 20, 0 ≤ *d* ≤ 11, *sn =* 6.4
  - *r =* 21, 0 ≤ *d* ≤ 11, *sn =* 6.3
  - *r =* 26, 0 ≤ *d* ≤ 11, *sn =* 5.8
  - *r =* 15, *d =* 6, 6.5 ≤ *sn* ≤ 7.5
  - *r =* 16, *d =* 6, 5.8 ≤ *sn* ≤ 7.8
  - *r =* 21, *d =* 6, 5.8 ≤ *sn* ≤ 6.7
  - *r =* 24, *d =* 6, 5.6 ≤ *sn* ≤ 6.8
  - *r =* 26, *d =* 6, 5.4 ≤ *sn* ≤ 6.2
- For the “movement” period dataset (*z* is fixed to 3; *sn* is varied in steps of 0.1, unless noted otherwise):
  - *r =* 16, *d =* 4, 7 ≤ *sn* ≤ 8 (varied in steps of 0.2)
  - *r =* 18, *d =* 4, 6.5 ≤ *sn* ≤ 7.5 (varied in steps of 0.2)
  - *r =* 20, *d =* 4, 6 ≤ *sn* ≤ 7 (varied in steps of 0.2)
  - *r =* 22, *d =* 5, 5.9 ≤ *sn* ≤ 6.5
  - *r =* 24, *d =* 5, 5.76 ≤ *sn* ≤ 6.36
  - *r =* 26, *d =* 5, 5.63 ≤ *sn* ≤ 6.23
  - *r =* 28, *d =* 5, 5.5 ≤ *sn* ≤ 6.1
  - *r =* 30, *d =* 6, 5.4 ≤ *sn* ≤ 5.9
  - *r =* 32, *d =* 6, 5.3 ≤ *sn* ≤ 5.8
  - *r =* 34, *d =* 6, 5.2 ≤ *sn* ≤ 5.7
  - *r =* 36, *d =* 6, 5.1 ≤ *sn* ≤ 5.6

For the comparison of the computed stationary or dynamic stationary quality to the quality estimate based on non-stationary simulasions (Figs. 7 and 8), we collected signatures in the following way. We performed 1,000 non-stationary simulations for each of the two datasets (“wait” and “movement” periods) and extracted all PRs of size 3. For each of these PRs, we derived its quality estimate from the quality spectrum (obtained from a separate set of 10.000 non-stationary simulations), and stored the quality value along with the corresponding signature parameters (size, repetition, duration, and pattern novelty). From these collections, we randomly picked up at most 10 signatures per quality bin of width 0.1 in the range [1, 4], resulting in 264 signatures for the “wait” period dataset and 267 signatures for the “movement” period dataset.

## Footnotes

1 This article is published as part of the Special Issue onNeuronal Spike Timing: Precision, Reliability, and theNeural Code from 1995-2025

